# Levosimendan Ameliorates Adverse Pulmonary Vascular Remodeling in Group-2 Pulmonary Hypertension

**DOI:** 10.1101/2025.10.23.684000

**Authors:** Antonella Abi Sleimen, Ruaa Al-Qazazi, Yann Grobs, Yahe Xu, Tsukasa Shimauchi, Ashley Martin, Danchen Wu, Reem El Kabbout, Manon Mougin, Charlie Théberge, Steeve Provencher, Olivier Boucherat, Sébastien Bonnet, Stephen L Archer, François Potus

**Affiliations:** Centre de Recherche de l’Institut Universitaire de Cardiologie et de Pneumologie de Québec (CRIUCPQ), Québec, Québec, Canada; Department of Medicine, Queens University, Kingston, Ontario, Canada; Institute for Lung Health (ILH), German Center for Lung Research (DZL) Cardio-Pulmonary Institute (CPI) Justus-Liebig University Giessen, Giessen, Germany

## Abstract

**Aims:** Pulmonary hypertension (PH) due to left heart disease (Group-2PH) is the most common form of PH and comprises two distinct subtypes: isolated post-capillary-PH (IpcPH) and combined post-and pre-capillary-PH (CpcPH). Despite its high prevalence and poor prognosis, no targeted therapies are currently approved, largely due to the absence of reliable preclinical models that recapitulate these human hemodynamic phenotypes. Levosimendan, a calcium sensitizer with inotropic and vasodilatory properties, has shown promise in early clinical trials for Group-2PH, but its mechanisms of action remain unclear. This study aimed to develop and validate experimental models of IpcPH and CpcPH and to assess the therapeutic effects of levosimendan on pulmonary vascular remodeling, inflammation, and cardiac function to support ongoing clinical translation.

**Methods and Results:** In a multicentre preclinical study, we established two rodent models that faithfully replicate the human IpcPH and CpcPH hemodynamic profiles. CpcPH animals exhibited severe pulmonary vascular remodeling, inflammatory cell infiltration, and a distinct pro-proliferative transcriptomic signature, whereas IpcPH animals showed minimal pulmonary vascular involvement. Levosimendan (3 mg/kg/day, 3 weeks) improved biventricular function and pulmonary hemodynamics in both models. In CpcPH, levosimendan additionally reduced pulmonary vascular remodeling, attenuated inflammation, and partially reversed disease-associated transcriptomic reprogramming. Transcription factor enrichment analysis identified NF-κB as a key upstream regulator inhibited by treatment. In a translational extension, nine circulating inflammation-related-proteins differentiated CpcPH from IpcPH patients; among them, TNF, IL-12B, 4E-BP1, NT-3, NGF, FGF21, and FGF23 predicted poor survival. IL-18 and 4E-BP1 were elevated in CpcPH lungs and decreased following levosimendan treatment.

**Conclusions:** Inflammation is a major contributor to adverse pulmonary vascular remodeling in CpcPH. Levosimendan improves cardiac performance and mitigates pulmonary vascular inflammation and remodeling, supporting its potential as a dual-action therapeutic agent in Group-2PH. These findings validate novel preclinical models and provide mechanistic evidence reinforcing ongoing clinical evaluation of levosimendan in this condition.

**Translational perspective:** Group-2 PH lacks targeted therapies, partly due to the absence of validated preclinical models. We validated models recapitulating human IpcPH and CpcPH and identified inflammation as a key driver of pulmonary vascular remodeling in CpcPH. Levosimendan improved biventricular function and reduced vascular remodeling and inflammation through NF-κB inhibition. Circulating IL-18 and 4E-BP1 reflected disease severity and treatment response. These findings establish robust translational models, reveal inflammatory mechanisms underlying CpcPH, and provide mechanistic evidence supporting ongoing clinical trials of levosimendan as a dual-action therapeutic strategy in Group-2 PH.

## Introduction

Group-2 pulmonary hypertension (PH), also known as PH due to left heart disease (PH-LHD), is the most common subtype of PH, with an estimated prevalence of 67.4 per 100,000 and accounting for 60–70% of cases in population-based studies(1–3). Despite its prevalence, Group-2PH remains one of the most lethal forms of PH, with a 5-year survival of only ∼30%(1). Clinically, it is classified into two subtypes: isolated post-capillary PH (IpcPH), driven by elevated left atrial pressure, and combined post-and pre-capillary PH (CpcPH), which involves pulmonary arterial remodeling and increased pulmonary vascular resistance. Both forms meet the diagnostic thresholds of mean pulmonary arterial pressure (mPAP) > 20 mmHg and pulmonary artery wedge pressure (PAWP) > 15 mmHg. However, IpcPH is characterized by pulmonary vascular resistance (PVR) < 3 Wood Units (WU), while CpcPH presents with PVR > 3 WU and more extensive vascular changes(4,5). Histological studies confirm that CpcPH, but not IpcPH, is associated with significant pulmonary vascular remodeling(4,5), which translates clinically into a 2.5-fold higher mortality risk in CpcPH patients compared to those with IpcPH(6).

Current therapeutic strategies for Group-2 PH focus on treating underlying left heart disease. Sodium–glucose co-transporter 2 inhibitors (SGLT2i) have demonstrated rapid pulmonary pressure reductions in heart failure with preserved ejection fraction (HFpEF) (7). However, this strategy has limited efficacy in CpcPH, where vascular remodeling and inflammation dominate. Several trials repurposing pulmonary arterial hypertension (PAH, Group-1 PH) therapies for Group-2 PH have failed to show benefit and, in some cases, have caused harm(8). Critically, no targeted therapies exist for either IpcPH or CpcPH, partly due to the prevailing approach of treating Group-2 PH as a homogeneous entity. The absence of subtype-specific preclinical models has further hindered mechanistic discoveries and therapeutic advances.

Levosimendan is a calcium sensitizer with inotropic and vasodilatory properties, approved in Europe and other regions for treating decompensated heart failure. It exerts positive effects by enhancing cardiac contractility and opening ATP-sensitive potassium channels, leading to systemic vasodilation(9). In patients with heart failure with reduced ejection fraction (HFrEF), levosimendan has shown favorable hemodynamic effects and improvements in right ventricular (RV) function(10,11) (12–14). It is currently under investigation in clinical trials for Group-2 PH. A phase 2 trial (NCT03541603) showed that once-weekly levosimendan infusions reduced PAWP and improved 6-minute walk distance in PH-HFpEF patients. Ongoing trials include phase 3 studies further evaluating its efficacy (NCT03624010, NCT05983250) (15).

Preclinical studies in PAH (Group-1 PH) suggest that levosimendan may mitigate pulmonary vascular remodeling via inhibition of smooth muscle cell proliferation, improved endothelial function, and reduced inflammation(16,17). Ex vivo, levosimendan decreases segmental pulmonary vascular resistance and relaxes human pulmonary vessels in precision-cut lung slices(18). However, its effects on IpcPH and CpcPH remain unexplored.

To address this gap, we aimed to evaluate the effects of levosimendan on pulmonary vascular remodeling and hemodynamics in validated preclinical models of IpcPH and CpcPH. Using clinical hemodynamic definitions, we characterized rodent models that recapitulate the distinct features of both subtypes. We then leveraged a well-characterized human cohort for translational validation. We found that levosimendan markedly reduced pulmonary vascular remodeling, inflammation, and transcriptomic reprogramming in CpcPH animals but had limited effects in IpcPH. Furthermore, circulating inflammatory markers differentiated IpcPH from CpcPH patients. Two cytokines—IL-18 and 4E-BP1—were upregulated in CpcPH lungs and downregulated following treatment. These results highlight levosimendan’s dual therapeutic action and underscore the importance of inflammation in CpcPH pathogenesis and treatment response.

## Methods

For the purpose of replicating procedures or reproducing results, the data, analytic methods, and study materials can be made available upon request to the corresponding author, who oversees the information. All experiments were conducted in compliance with the biosafety and human (CER-20735) and animal (CPAUL-2022-1049) ethical guidelines of Laval University, as well as the biosafety and animal ethical guidelines of Queen’s University (#2308). Informed consent was obtained from all patients before the study began. All experiments were conducted and analyzed blindly, following the guidelines outlined in(19).

### Animal Model and Experimental Design

#### Model 1 – Isolated Post Capillary Pulmonary Hypertension at Queen’s University

Following one week of acclimation, male Sprague Dawley rats (30 animals, Charles River, weighing 200±10 g; aged 42±4 days) underwent supracoronary aortic banding (SAB) surgery as outlined in(20), while 5 animals received sham surgery. Seven weeks post-surgery, SAB animals were randomly assigned to two groups: 15 animals received Levosimendan treatment (3mg/kg/day via oral gavage, Sigma, L5545), and 15 animals received vehicle treatment for a period of three weeks. Pulmonary hypertension, pulmonary vascular remodeling, and pulmonary hemodynamics were assessed at the beginning of the treatment and at the end of the ninth week for all groups (**Figure S1 A**).

#### Model 2 – Combined Postcapillary and Precapillary Pulmonary Hypertension at Quebec

CpcPH was induced by a combination of SAB surgery and metabolic syndrome (MetS), as outlined in(21). Briefly, after an acclimatization period of one week, 30 rats underwent SAB surgery, while 5 rats received sham surgery. MetS was subsequently induced in all animals by either a high-fat diet (Research Diets; D12492, 60% kcal% fat ad libitum) or olanzapine (Eli Lilly; 4 mg/kg body weight, intraperitoneal injection every two days) for a duration of nine weeks. Seven weeks post-surgery, animals with SAB + MetS were randomly assigned to two groups: 15 animals received Levosimendan treatment (3mg/kg/day via oral gavage, Sigma, L5545), and 15 animals received vehicle treatment for a period of three weeks. Pulmonary hypertension and pulmonary vascular adaptation were evaluated at the onset of treatment and at the conclusion of the ninth week for all groups. The success rate of surgery and obtaining hemodynamic parameters across all protocols was 80%. While our research involved multi-centric animal models, data analysis, histology, and transcriptomic studies were centralized in Quebec, a single center. This approach was taken to ensure consistency and improve quality control (**Figure S1 B**).

### Haemodynamic

#### Echocardiography, Right and Left Heart Catheterization

Transthoracic echocardiography was performed using a Vevo® 2100 (FUJIFILM VisualSonics Inc.) was performed as described in(21) and supplemental. Invasive, closed-chest, right heart and left heart catheterization were performed using the Transonic Scisense ADV500 Pressure-Volume Measurement System and analyzed with Labscibe2 software from Work System, Inc, following previously described methods(21,22) and supplemental.

***Histology and biochemistry*** (See supplemental)

### RNA-sequencing

RNA extraction was performed from lung tissue samples from model 1 (5 Sham, 5 SAB (IpcPH), 5 SAB (IpcPH) + Levosimendan) and model 2 (5 Sham-Mets, 5 SAB-Mets (CpcPH), 5 SAB-Mets (CpcPH) + Levosimendan) using the Direct-zol RNA Miniprep Kit (#R2051, Zymo Research, USA). Subsequently, RNA libraries were constructed using the Illumina Stranded mRNA Library Preparation Kit. The samples were sequenced on an Illumina NovaSeq platform, generating 150bp paired-end reads with a target of 50 million clusters per sample (100 million reads total). As a cutoff criterion for the study, transcripts with an adjusted p-value < 0.01 and a log2 fold change > 1 were included. Gene ontology analysis was conducted using KEGG, Gene Ontology Biological Function, Gene Ontology Cellular Component, and Gene Ontology Molecular Function, and TF.Target.Tfact databases, employing ShinnyGo(23). The datasets generated and/or analyzed during the current study are available in the GEO repository (accession number pending). Cellular deconvolution was conducted using CIBERSORTx(24), leveraging a single-cell dataset derived from PAH rat lungs(25) as the reference signature matrix.

### Olink Proteomic Analysis

Peripheral blood samples were collected from each participant in EDTA tubes via the central venous line at the time of right heart catheterization. Samples were immediately processed by centrifugation to isolate plasma, which was then stored at −80 °C until proteomic analysis. Baseline clinical data were extracted from institutional medical records. Plasma samples from 88 patients were analyzed using the Olink® Target 96 Inflammation panel (Olink Proteomics, Uppsala, Sweden), a proximity extension assay designed for high-throughput multiplex quantification of 96 immune-related proteins, including cytokines and chemokines. The assay relies on dual antibody binding to target epitopes, with each antibody conjugated to a unique oligonucleotide. Upon simultaneous binding, the oligonucleotides hybridize, allowing subsequent amplification and quantification by next-generation sequencing. Protein expression levels are reported as log2-normalized protein expression (NPX) values. All assays were performed at the Olink-certified core facility at the Lunenfeld-Tanenbaum Research Institute (Toronto, ON, Canada). Laboratory personnel were blinded to clinical diagnosis and group allocation.

## Statistical Analysis

For preclinical data, values are presented as mean ± standard error of the mean (SEM) and were analyzed using one-way ANOVA followed by Tukey’s multiple comparisons test or Kruskal–Wallis test followed by Dunn’s post hoc test, depending on data distribution. Human data are reported as mean ± standard deviation for normally distributed variables and as median with interquartile range [IQR] for non-normally distributed variables, unless otherwise specified. Categorical variables were compared using chi-square tests with Yates’ continuity correction or Fisher’s exact test, as appropriate. Ordinal variables were analyzed using the Mann–Whitney U test. Continuous variables were compared using Student’s t-test, Welch’s t-test, or Mann–Whitney U test, based on distribution and variance homogeneity. Correlations between circulating protein levels and clinical parameters (e.g., pulmonary vascular resistance, cardiac index, stroke volume index, NT-proBNP) were assessed using Spearman’s rank correlation. Correlation plots were generated using the ggplot2 package in R. Receiver operating characteristic (ROC) curve analyses were performed with the pROC package to evaluate the discriminatory performance of candidate plasma proteins for distinguishing CpcPH from IpcPH. To identify a minimal predictive variable set for Group 2 PH classification, we conducted LASSO-penalized logistic regression using the glmnet package in R. The model incorporated demographics, comorbidities, and protein levels. Patients with missing data were excluded. A design matrix was created with model.matrix(), and the outcome variable was defined as CpcPH = 1 and IpcPH = 0. Ten-fold cross-validation determined the optimal penalty parameter (λ), and predictors with non-zero coefficients at lambda.min were retained. LASSO-derived scores and predicted probabilities were computed, and model performance assessed via ROC analysis. Prognostic evaluation of differentially expressed proteins identified by the Olink assay was conducted using univariate and multivariate Cox proportional hazards models (survival and ggplot2 packages). The primary endpoint was all-cause mortality, with time-to-event defined from the time of blood sampling to death or last follow-up. Median follow-up was estimated both as a simple median for surviving patients (1.79 years) and using the reverse Kaplan–Meier method (5.77 years). Additional Cox models assessed the incremental prognostic value of individual biomarkers and the composite LASSO score when added to a base model including log(NT-proBNP). Model performance was evaluated using C-statistics, Integrated Discrimination Improvement (IDI), and Net Reclassification Index (NRI). Kaplan–Meier curves were generated using ROC-optimized cut-off values to dichotomize patients into “high” and “low” expression groups. Time was calculated from right heart catheterization to death or last follow-up, with risk tables included. Survival analyses were performed with the survival and survminer packages. All statistical tests were two-sided, and p < 0.05 was considered significant. Analyses were conducted using R version 4.3.2 (R Foundation, Vienna), GraphPad Prism 9.5.0 (GraphPad Software, CA), and SPSS version 25 (IBM, NY).

## Results

### Characterization of IpcPH and CpcPH Preclinical Models

Prior to commencing our investigation, we conducted a detailed characterization of two preclinical models representative of Group 2 PH, adhering to the hemodynamic criteria that distinguish IpcPH and CpcPH as observed in clinical scenarios. These criteria include mPAP > 20 mmHg, PAWP > 15 mmHg, PVR ≤ WU, TPG ≤ 12 mmHg for IpcPH, and mPAP > 20 mmHg, PAWP > 15 mmHg, PVR > 3 WU, and TPG > 12 mmHg for CpcPH (**Table 1**). In our experimental setting, Left Ventricular End-Diastolic Pressure (LVEDP) was used instead of PAWP, as it is a more direct measurement of left heart filling pressures. The translation of the PVR threshold to our rat models was adjusted to account for their significantly lower cardiac output compared to humans. The SAB surgery in model-1 elicited a marked elevation in mPAP and LVEDP when compared with Sham controls, yet PVR and TPG remained unchanged. In contrast, model-2, encompassing SAB + Mets, demonstrated substantial increases across mPAP, LVEDP, PVR, and TPG compared to Sham + Mets controls. Furthermore, the comparison between SAB and SAB + Mets groups revealed that the SAB + Mets group manifested significantly higher mPAP, PVR, and TPG (**Table 1**). To ensure the observed hemodynamic disparities were not artifacts of technical variability in the severity of aortic constriction, we assessed the systolic pressure gradient before and after banding. The absence of a significant difference in Trans Banding Gradient (TBG) between the SAB groups of both models corroborates the technical equivalency of these models. Thus, the hemodynamic profile in model-1 aligns with the clinical definition of IpcPH, whereas model-2 aligns with CpcPH. Hereafter in our manuscript, these will be referred to as IpcPH and CpcPH, respectively.

**Table 1:**
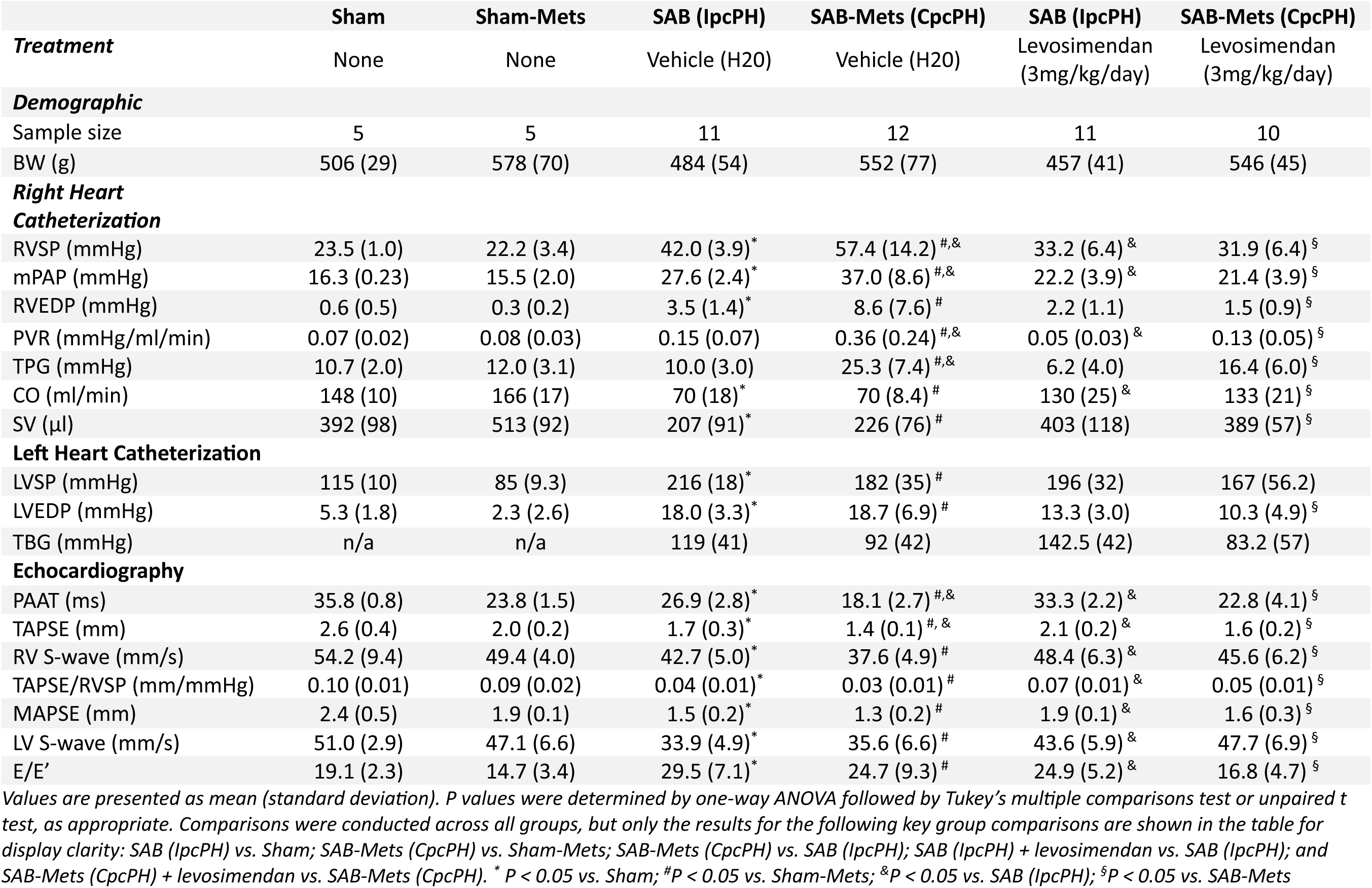
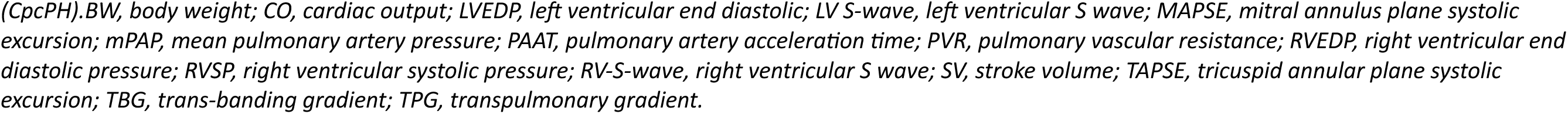
Preclinical Model haemodynamic characterisation.

|  | Sham | Sham-Mets | SAB (IpcPH) | SAB-Mets (CpcPH) | SAB (IpcPH) | SAB-Mets (CpcPH) |
| --- | --- | --- | --- | --- | --- | --- |
| **Treatment** | None | None | Vehicle (H2O) | Vehicle (H2O) | Levosimendan (3mg/kg/day) | Levosimendan (3mg/kg/day) |
| **Demographic** |  |  |  |  |  |  |
| Sample size | 5 | 5 | 11 | 12 | 11 | 10 |
| BW (g) | 506 (29) | 578 (70) | 484 (54) | 552 (77) | 457 (41) | 546 (45) |
| **Right Heart Catheterization** |  |  |  |  |  |  |
| RVSP (mmHg) | 23.5 (1.0) | 22.2 (3.4) | 42.0 (3.9)* | 57.4 (14.2)<sup>#,&</sup> | 33.2 (6.4)<sup>&</sup> | 31.9 (6.4)<sup>§</sup> |
| mPAP (mmHg) | 16.3 (0.23) | 15.5 (2.0) | 27.6 (2.4)* | 37.0 (8.6)<sup>#,&</sup> | 22.2 (3.9)<sup>&</sup> | 21.4 (3.9)<sup>§</sup> |
| RVEDP (mmHg) | 0.6 (0.5) | 0.3 (0.2) | 3.5 (1.4)* | 8.6 (7.6)<sup>#</sup> | 2.2 (1.1) | 1.5 (0.9)<sup>§</sup> |
| PVR (mmHg/ml/min) | 0.07 (0.02) | 0.08 (0.03) | 0.15 (0.07) | 0.36 (0.24)<sup>#,&</sup> | 0.05 (0.03)<sup>&</sup> | 0.13 (0.05)<sup>§</sup> |
| TPG (mmHg) | 10.7 (2.0) | 12.0 (3.1) | 10.0 (3.0) | 25.3 (7.4)<sup>#,&</sup> | 6.2 (4.0) | 16.4 (6.0)<sup>§</sup> |
| CO (ml/min) | 148 (10) | 166 (17) | 70 (18)* | 70 (8.4)<sup>#</sup> | 130 (25)<sup>&</sup> | 133 (21)<sup>§</sup> |
| SV (μl) | 392 (98) | 513 (92) | 207 (91)* | 226 (76)<sup>#</sup> | 403 (118) | 389 (57)<sup>§</sup> |
| **Left Heart Catheterization** |  |  |  |  |  |  |
| LVSP (mmHg) | 115 (10) | 85 (9.3) | 216 (18)* | 182 (35)<sup>#</sup> | 196 (32) | 167 (56.2) |
| LVEDP (mmHg) | 5.3 (1.8) | 2.3 (2.6) | 18.0 (3.3)* | 18.7 (6.9)<sup>#</sup> | 13.3 (3.0) | 10.3 (4.9)<sup>§</sup> |
| TBG (mmHg) | n/a | n/a | 119 (41) | 92 (42) | 142.5 (42) | 83.2 (57) |
| **Echocardiography** |  |  |  |  |  |  |
| PAAT (ms) | 35.8 (0.8) | 23.8 (1.5) | 26.9 (2.8)* | 18.1 (2.7)<sup>#,&</sup> | 33.3 (2.2)<sup>&</sup> | 22.8 (4.1)<sup>§</sup> |
| TAPSE (mm) | 2.6 (0.4) | 2.0 (0.2) | 1.7 (0.3)* | 1.4 (0.1)<sup>#,&</sup> | 2.1 (0.2)<sup>&</sup> | 1.6 (0.2)<sup>§</sup> |
| RV S-wave (mm/s) | 54.2 (9.4) | 49.4 (4.0) | 42.7 (5.0)* | 37.6 (4.9)<sup>#</sup> | 48.4 (6.3)<sup>&</sup> | 45.6 (6.2)<sup>§</sup> |
| TAPSE/RVSP (mm/mmHg) | 0.10 (0.01) | 0.09 (0.02) | 0.04 (0.01)* | 0.03 (0.01)<sup>#</sup> | 0.07 (0.01)<sup>&</sup> | 0.05 (0.01)<sup>§</sup> |
| MAPSE (mm) | 2.4 (0.5) | 1.9 (0.1) | 1.5 (0.2)* | 1.3 (0.2)<sup>#</sup> | 1.9 (0.1)<sup>&</sup> | 1.6 (0.3)<sup>§</sup> |
| LV S-wave (mm/s) | 51.0 (2.9) | 47.1 (6.6) | 33.9 (4.9)* | 35.6 (6.6)<sup>#</sup> | 43.6 (5.9)<sup>&</sup> | 47.7 (6.9)<sup>§</sup> |
| E/E' | 19.1 (2.3) | 14.7 (3.4) | 29.5 (7.1)* | 24.7 (9.3)<sup>#</sup> | 24.9 (5.2)<sup>&</sup> | 16.8 (4.7)<sup>§</sup> |

Values are presented as mean (standard deviation). *P* values were determined by one-way ANOVA followed by Tukey's multiple comparisons test or unpaired *t* test, as appropriate. Comparisons were conducted across all groups, but only the results for the following key group comparisons are shown in the table for display clarity: SAB (IpcPH) vs. Sham; SAB-Mets (CpcPH) vs. Sham-Mets; SAB-Mets (CpcPH) vs. SAB (IpcPH); SAB (IpcPH) + levosimendan vs. SAB (IpcPH); and SAB-Mets (CpcPH) + levosimendan vs. SAB-Mets (CpcPH). \* *P* < 0.05 vs. Sham; <sup>#</sup>*P* < 0.05 vs. Sham-Mets; <sup>&</sup>*P* < 0.05 vs. SAB (IpcPH); <sup>§</sup>*P* < 0.05 vs. SAB-Mets
*(CpcPH).BW, body weight; CO, cardiac output; LVEDP, left ventricular end diastolic; LV S-wave, left ventricular S wave; MAPSE, mitral annulus plane systolic excursion; mPAP, mean pulmonary artery pressure; PAAT, pulmonary artery acceleration time; PVR, pulmonary vascular resistance; RVEDP, right ventricular end diastolic pressure; RVSP, right ventricular systolic pressure; RV-S-wave, right ventricular S wave; SV, stroke volume; TAPSE, tricuspid annular plane systolic excursion; TBG, trans-banding gradient; TPG, transpulmonary gradient.*

Subsequent comprehensive characterization of pulmonary, right and left ventricular, as well as cardiovascular hemodynamic parameters in Sham, IpcPH, and CpcPH cohorts revealed expected deviations. Relative to their respective Sham counterparts, both IpcPH and CpcPH animals demonstrated significant decrements in left ventricular function, as evidenced by increased Left Ventricular Systolic Pressure (LVSP), augmented E/E’ ratio, and diminished Mitral Annular Plane Systolic Excursion (MAPSE) and LV S’ wave (**Table 1**). Right ventricular function was similarly compromised, indicated by reduced Right Ventricular End-Diastolic Pressure (RVEDP), Tricuspid

Annular Plane Systolic Excursion (TAPSE), RV S’ wave, and a lower TAPSE/RV Systolic Pressure (RVSP) coupling ratio. Pulmonary hemodynamics were adversely affected, denoted by heightened RVSP, diminished Pulmonary Artery Acceleration Time (PAAT), and overall cardiac function was impaired, as shown by a decrease in Cardiac Output (CO) and Stroke Volume (SV). Most notably, the CpcPH model exhibited significantly more severe pulmonary hemodynamic impairment than the IpcPH model, with higher mPAP (37.0 ± 8.6 v.s. 37.6 ± 2.4 mmHg), RVSP (57.4 ± 14.2 v.s. 42.0 ± 3.9 mmHg), PVR (0.36 ± 0.24 v.s. 0.15 ± 0.07 mmHg/ml/min), and TPG (25.3 ± 7.4 v.s. 10.0 ± 3.0 mmHg), alongside lower PAAT (18.1 ± 2.7 v.s. 26.9 ± 2.8 ms) (**Table 1**). These observations indicate a markedly exacerbated impact on lung hemodynamics in conditions of CpcPH, which could denote a more extensive degree of pulmonary vascular remodeling compared to IpcPH animals.

To evaluate the translational relevance of our preclinical models, we conducted a comprehensive clinical and hemodynamic characterization of patients with IpcPH and CpcPH (**Table 2**). As expected, CpcPH patients exhibited more severe pulmonary hemodynamics compared to IpcPH, including higher mPAP, sPAP, PVR, TPG, along with reduced pulmonary arterial compliance. CpcPH patients also trended to have greater pulmonary dysfunction, as indicated by lower mixed venous oxygen saturation (SvO₂), peripheral oxygen saturation (SpO₂), Forced Expiratory Volume in 1 second (FEV1), and diffusing capacity of the lungs for carbon monoxide (DLCO) (**Table 2**). Although left ventricular shortening fraction was similar between groups, CpcPH patients had more pronounced RV dysfunction, RV dilation, and reduced cardiac index, along with higher circulating NT-proBNP levels (**Table 2**). No significant differences were observed between groups in terms of comorbidities or medication use. Consistent with prior reports, CpcPH patients had significantly worse outcomes, with a 7-year survival rate of 21.1% compared to 60.6% in IpcPH patients, and a median survival of 3.7 years versus 8.7 years, respectively (**Figure S2 A**). Although no risk score has been specifically validated for Group-2 PH, we applied the 2022 ESC/ERS risk stratification algorithm developed for Group-1 PH. In line with survival data, CpcPH patients exhibited higher risk scores (**Table 2**).

**Table 2.** Clinical characteristics of patients with isolated post-capillary PH (IpcPH) and combined post-and pre-capillary PH (CpcPH).

|  | IpcPH (n=38) | CpcPH (n=52) | P Value |
| --- | --- | --- | --- |
| Demographics |  |  |  |
| Male | 21 (55.3%) | 19 (36.5%) | 0.1209 |
| Age (years) | 70.50 (67.00-75.00) | 72.50 (66.75-77.25) | 0.1720 |
| Caucasian | 38 (100%) | 52 (100%) |  |
| Size (meter) | 1.63 (1.59-1.71) | 1.58 (1.53-1.66) | 0.0443 |
| Weight (kg) | 76.55 (66.28-94.00) | 76.00 (66.25-87.85) | 0.5982 |
| BMI (kg/m<sup>2</sup>) | 30.06 ± 6.33 | 29.93 ± 5.22 | 0.9184 |
| Smoking |  |  |  |
| Non-smoker | 13 (34.2%) | 23 (44.2%) | 0.2251 |
| Former-smoker | 21 (55.3%) | 27 (51.9%) |  |
| Current-smoker | 4 (10.5%) | 2 (3.8%) |  |
| Comorbidities |  |  |  |
| Diabetes | 11 (28.9%) | 25 (48.1%) | 0.1070 |
| Dyslipidemia | 22 (57.9%) | 26 (50.0%) | 0.5978 |
| Hypertension | 24 (63.2%) | 35 (67.3%) | 0.8535 |
| Obesity | 17 (44.7%) | 28 (53.8%) | 0.5220 |
| CTDs* | 6 (15.8%) | 2 (3.8%) | 0.0660 |
| Medications |  |  |  |
| Diuretics | 38 (100%) | 52 (100%) |  |
| Beta-blockers | 20 (52.6%) | 34 (65.4%) | 0.3164 |
| CCBs | 15 (39.5%) | 20 (38.5%) | 0.9032 |
| ERAs | 4 (10.5%) | 6 (11.5%) | >0.9999 |
| Antidiabetics | 12 (31.6%) | 20 (38.5%) | 0.6521 |
| Statins | 22 (57.9%) | 27 (51.9%) | 0.7282 |
| Functional Parameters |  |  |  |
| WHO Class |  |  |  |
| Class I | 1 (2.6%) | 2 (3.9%) | 0.1635 |
| Class II | 17 (44.7%) | 15 (29.4%) |  |
| Class III | 19 (50.0%) | 30 (58.9%) |  |
| *Class IV* | 1 (2.6%) | 4 (7.8%) |  |
| **6 Minute Walking Distance (meter)** | 358.04 ± 127.63 | 256.59 ± 104.51 | 0.0010* |
| **2022 ESC/ERS Risk Score** | 2.00 (1.63-2.22) | 2.30 (2.00-2.70) | <0.0001* |
| **2022 ESC/ERS Risk Stratification** |  |  |  |
| *Low Risk* | 6 (15.8%) | 0 |  |
| *Intermediate Risk* | 28 (73.7%) | 32 (61.5%) | 0.0002* |
| *High Risk* | 4 (10.5%) | 20 (38.5%) |  |
| **Abnormality in V/Q Scan** | 6 (33.3%) | 17 (54.8%) | 0.2472 |
| **SpO2 (%)** | 97.00 (95.50-98.00) | 96.00 (91.00-98.00) | 0.0810 |
| **Right heart catheterization** |  |  |  |
| *mRAP (mmHg)* | 9.00 (6.00-14.00) | 10.00 (8.75-15.25) | 0.1278 |
| *sPAP (mmHg)* | 45.71 ± 8.66 | 70.38 ± 17.34 | <0.0001* |
| *dPAP (mmHg)* | 17.00 (14.25-22.75) | 25.00 (20.00-30.00) | <0.0001* |
| *mPAP (mmHg)* | 27.00 (24.00-32.75) | 41.00 (34.75-46.50) | <0.0001* |
| *sBP (mmHg)* | 145.53 ± 22.01 | 135.53 ± 23.22 | 0.0464* |
| *dBp (mmHg)* | 70.17 ± 12.42 | 70.04 ± 14.07 | 0.9653 |
| *mBP (mmHg)* | 88.39 ± 15.32 | 87.66 ± 13.81 | 0.8182 |
| *PAWP (mmHg)* | 18.11 ± 6.03 | 19.71 ± 6.07 | 0.2190 |
| *PVR (WU)* | 2.09 (1.53-2.46) | 4.86 (3.95-7.42) | <0.0001* |
| *TPG (mmHg)* | 10.00 (8.00-12.00) | 21.5 (17.00-26.25) | <0.0001* |
| *Cardiac Index (L/min/m2)* | 2.70 (2.19-3.16) | 2.33 (1.99-2.70) | 0.0018* |
| *SVI (mL/m2)* | 41.04 (35.71-48.48) | 31.74 (26.63-37.35) | <0.0001* |
| *SVI (L/m2)* | 0.04 (0.04-0.05) | 0.03 (0.03-0.04) | <0.0001* |
| *Pulmonary vascular compliance (mL/mmHg)* | 3.00±1.17 | 1.43±0.51 | <0.0001* |
| *SvO2 (%)* | 66.86 ± 7.67 | 61.50 ± 6.97 | 0.0009* |
| *Heart Rate (beat/min)* | 65.50 (60.00-76.75) | 73.00 (62.50-82.50) | 0.0960 |
| **Echocardiography** |  |  |  |
| *TAPSE (mm)* | 18.56 ± 3.99 | 17.14 ± 4.24 | 0.3720 |
| *TAPSE/sPAP* | 0.39 (0.31-0.52) | 0.25 (0.20-0.28) | 0.0010* |
| *LVEF (%)* | 60.00 (55.00-60.00) | 60.00 (55.00-60.00) | 0.4856 |
| **RV Function** |  |  |  |
| *Normal* | 22 (59.5%) | 16 (38.1%) | 0.0355* |
| Mildly Reduced | 9 (24.3%) | 11 (26.2%) |  |
| Moderately Reduced | 5 (13.5%) | 13 (31.0%) |  |
| Severely Reduced | 1 (2.7%) | 2 (4.8%) |  |
| Size of the Right Atrium |  |  |  |
| Normal | 13 (59.1%) | 8 (25.8%) | 0.0274* |
| Mildly Dilated | 0 | 4 (12.9%) |  |
| Moderately Dilated | 6 (27.3%) | 8 (25.8%) |  |
| Severely Dilated | 3 (13.6%) | 11 (35.5%) |  |
| Tricuspid Regurgitation Level |  |  |  |
| None 1/4 | 4 (11.4%) | 3 (6.5%) | 0.2306 |
| Mild 2/4 | 12 (34.3%) | 13 (28.3%) |  |
| Moderate 3/4 | 13 (37.1%) | 18 (39.1%) |  |
| Severe 4/4 | 6 (17.1%) | 12 (26.1%) |  |
| Pulmonary Function Test |  |  |  |
| FEV1 (L) | 1.98 ± 0.62 | 1.58 ± 0.47 | 0.0027* |
| FEV1/FVC (%) | 75.96 ± 8.21 | 72.71 ± 8.06 | 0.1010 |
| DLCO (%) | 74.00 (60.50-80.50) | 61.35 (50.25-78.75) | 0.0642 |
| Blood Tests |  |  |  |
| White Blood Cells (WBC, ×10<sup>9</sup>/L) | 7.68 ± 2.81 | 8.09 ± 2.58 | 0.3842 |
| Neutrophils (×10<sup>9</sup>/L) | 5.00 (3.65-6.37) | 5.35 (4.18-6.64) | 0.4069 |
| Lymphocytes (×10<sup>9</sup>/L) | 1.30 (1.00-1.58) | 1.10 (0.80-1.60) | 0.3618 |
| Neutrophil-to-Lymphocyte Ratio (NLR) | 3.87 (2.89-5.36) | 4.51 (2.74-7.72) | 0.4798 |
| Monocytes (×10<sup>9</sup>/L) | 0.60 (0.50-0.78) | 0.60 (0.47-0.80) | 0.8201 |
| Eosinophils (×10<sup>9</sup>/L) | 0.15 (0.10-0.20) | 0.10 (0.10-0.20) | 0.3307 |
| Basophils (×10<sup>9</sup>/L) | 0.00 (0.00-0.09) | 0.00 (0.00-0.10) | 0.8485 |
| Platelets (×10<sup>9</sup>/L) | 211.00 (175.00-242.00) | 189.00 (151.00-225.00) | 0.1997 |
| Hemoglobin (g/L) | 123.39 ± 17.19 | 123.79 ± 20.13 | 0.9227 |
| RDW (%) | 14.80 (13.85-15.90) | 15.75 (15.07-17.50) | 0.0018* |
| HbA1C | 0.06 ± 0.01 | 0.07 ± 0.01 | 0.0749 |
| NT-proBNP (pg/mL) | 743.00 (236.75-1797.00) | 1507.00 (937.00-3391.00) | 0.0008* |
| Creatinin (μM) | 95.50 (81.25-112.75) | 104.00 (71.50-135.50) | 0.4139 |
| eGFR (ml/min/1.73m<sup>2</sup>) | 63.81 ± 19.84 | 56.76 ± 26.20 | 0.1740 |
| *Triglycerides (mmol/L)* | 1.23 (0.98-1.41) | 1.58 (0.96-1.92) | 0.0659 |

Data are presented as mean ± standard deviation for normally distributed variables, and as median (interquartile range) for non-normally distributed variables. p values were calculated using the chi-square test with Yates' continuity correction or Fisher's exact test for categorical variables, the Mann–Whitney U test for ordinal variables, and the Student's t-test, Welch's t-test, or Mann–Whitney U test for numerical variables, depending on the distribution and variance homogeneity.
Sample sizes may vary due to missing data.
BMI, body mass index; CTDs, connective tissue diseases; CCBs, calcium channel blockers; ERAs, endothelin receptor antagonists; WHO Class, World Health Organization functional class; ESC/ERS, European Society of Cardiology/European Respiratory Society; V/Q scan, ventilation/perfusion lung scan; SpO<sub>2</sub>, peripheral oxygen saturation at rest; RHC, right heart catheterization; mRAP, mean right atrial pressure; sPAP/dPAP/mPAP, systolic/diastolic/mean pulmonary arterial pressure; sBP/DBP/mBP, systolic/diastolic/mean systemic blood pressure; PAWP, pulmonary arterial wedge pressure; PVR, pulmonary vascular resistance; TPG, transpulmonary gradient; CI, cardiac index; SVI, stroke volume index; SvO<sub>2</sub>, mixed venous oxygen saturation; TAPSE, tricuspid annular plane systolic excursion; sPAP, systolic pulmonary arterial pressure; LVEF, left ventricular ejection fraction; FEV<sub>1</sub>, forced expiratory volume in one second; FVC, forced vital capacity; DLCO, diffusing capacity of the lung for carbon monoxide; RDW, red cell distribution width; HBA1C, glycated hemoglobin; NT-proBNP, N-terminal pro B-type natriuretic peptide; eGFR, estimated glomerular filtration rate.
\*CTDs: Disseminated lupus erythematosus, Sjögren's syndrome, Rheumatoid Arthritis, Fibromyalgia, Takayasu arteritis, Scleroderma.

Overall, IpcPH and CpcPH preclinical models faithfully recapitulate the pulmonary vascular and functional impairments observed in human disease. However, unlike in patients, CpcPH animals did not exhibit significantly worse RV function than IpcPH rats, aside from a modest reduction in TAPSE, and showed no difference in global cardiac output compared to IpcPH animals.

### Increased Pulmonary Vascular Remodeling and Transcriptomic Reprogramming in CpcPH Animal Models

We conducted histological and transcriptomic analyses to determine whether the hemodynamic disparities between the CpcPH and IpcPH models were also evident at the cellular and molecular levels. Specifically, we assessed pulmonary vascular remodeling, PASMC proliferation, and apoptosis across Sham, IpcPH, Sham + Mets, and CpcPH groups. Consistent with our hemodynamic findings, CpcPH animals exhibited significantly increased pulmonary vascular remodeling, elevated PASMC proliferation, and reduced PASMC apoptosis compared to both IpcPH and sham controls (**Figure 1A–D**). While pulmonary vascular remodeling was increased in both IpcPH and CpcPH animals, relative to their respective sham groups (**Figure 1A, B**), the increase in PASMC proliferation and the decrease in apoptosis were uniquely observed in CpcPH animals (**Figure 1B, C**). Furthermore, lungs from CpcPH animals showed substantial infiltration of inflammatory cells, including macrophages (CD68+) and leukocytes (CD45+), compared to all other experimental conditions (IpcPH and sham controls) (**Figure 1A, E, F**).

**Figure 1.**
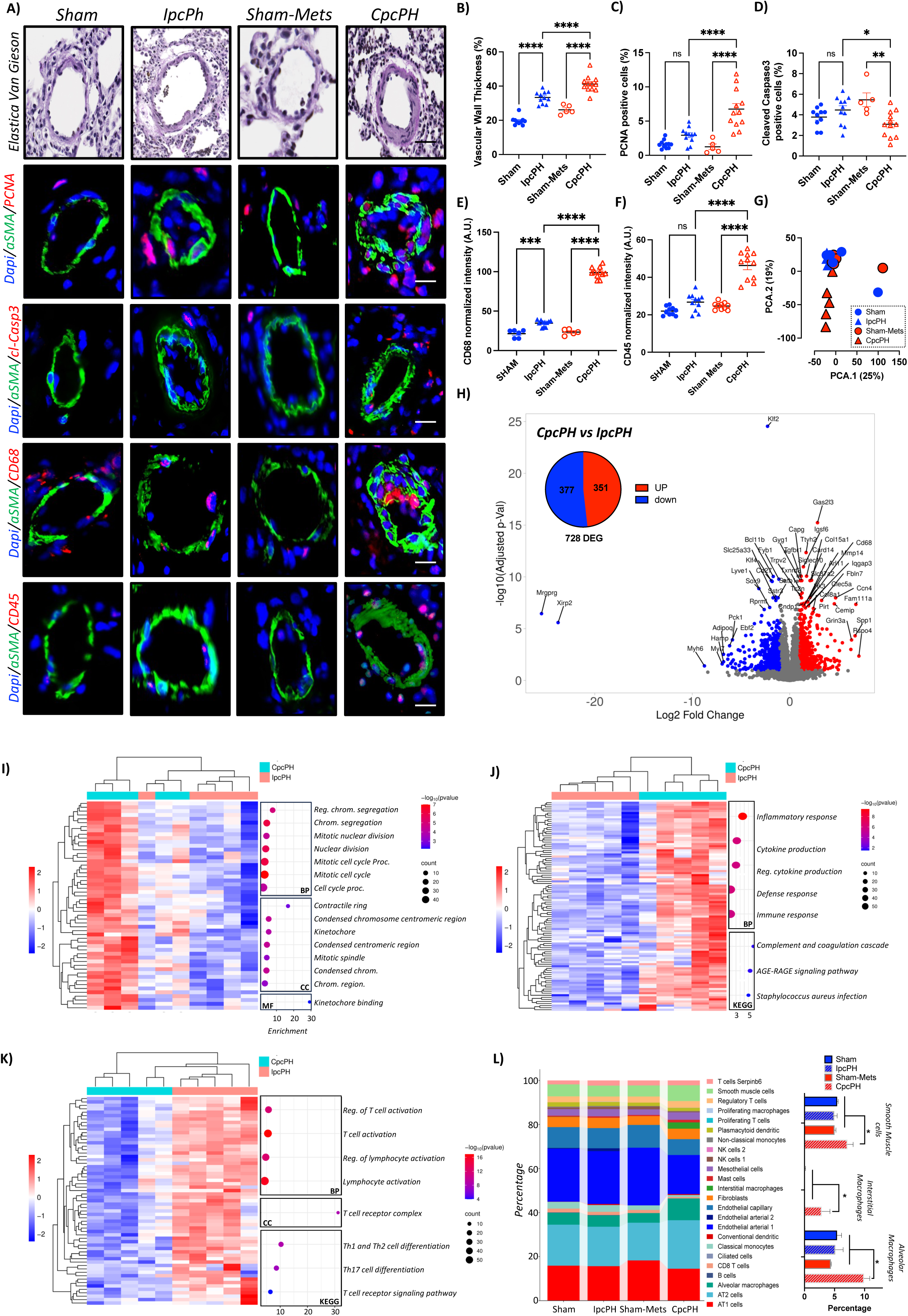
CpcPH preclinical model exhibits increased pulmonary vascular remodeling, inflammatory cell infiltration, and lung transcriptomic reprogramming. **A)** Representative histological and immunofluorescence images of lung sections from Sham, IpcPH, Sham-Mets, and CpcPH animals. Elastica Van Gieson staining shows increased medial thickening in CpcPH pulmonary arteries. Immunofluorescence staining reveals increased α-SMA⁺ PCNA⁺ proliferative PASMCs, reduced α-SMA⁺ cleaved caspase-3⁺ apoptotic cells, and increased monocyte (CD68⁺) and leukocyte (CD45⁺) cell infiltration in CpcPH lungs. Scale bars = 20 µm. **B**) Quantification of pulmonary vascular remodeling: medial thickness, **C**) percentage of PCNA⁺ PASMCs, **D**) cleaved caspase-3⁺ PASMCs, **E**) CD68 intensity, and **F**) CD45 intensity. n = 5–8/group. Data are mean ± SEM; ns = not significant, *p < 0.05, **p < 0.01, ****p < 0.0001. **G**) Principal component analysis (PCA) of lung transcriptomes reveals distinct clustering of CpcPH samples. **H**) Volcano plot of differentially expressed genes (DEGs) between CpcPH and IpcPH (log2FC > 1, adj. p < 0.01). 351 genes were upregulated and 377 downregulated in CpcPH lungs. **I, J**) Heatmaps and gene ontology analysis of upregulated DEGs in CpcPH. **I**) Enrichment in proliferation-associated pathways (e.g., chromosome segregation, mitotic cell cycle). **J**) Enrichment in inflammatory and cytokine signaling pathways. **K**) Downregulated DEGs are associated with lymphocyte activation and T cell differentiation. **L**) Cellular deconvolution analysis of lung transcriptomes using CIBERSORTx, showing increased macrophage populations (interstitial and alveolar) and altered immune cell composition in CpcPH compared to other groups.

Upon examining transcriptomic alterations in lung tissues, we executed an unsupervised Principal Component Analysis (PCA) which delineated a distinct clustering pattern. Specifically, Sham, Sham + Mets, and IpcPH samples shared a similar transcriptional profile, grouping closely in the PCA plot, while CpcPH samples formed a separate cluster, underscoring a distinct transcriptomic landscape for this condition (**Figure 1G**). Further investigation into differentially expressed genes (DEGs) reinforced this distinction, revealing that CpcPH lungs exhibited a significant differential expression profile compared to IpcPH (**Figure 1H**). In line with the histological findings, gene ontology analyses of the 351 upregulated DEGs in CpcPH highlighted their strong association with cellular proliferation pathways, such as chromosome segregation, nuclear division, and key mitotic components like the spindle and kinetochore binding (**Figure 1I**), as well as inflammatory processes, including inflammatory responses and cytokine production (**Figure 1J**). Intriguingly, the 377 downregulated DEGs were predominantly linked to lymphocyte activation and differentiation processes (**Figure 1K**), suggesting an immunomodulatory shift toward suppressed adaptive immune activity in CpcPH. We then performed cellular deconvolution using a publicly available single-cell dataset from PAH rat lungs(25) and observed an enrichment of macrophages (both interstitial and alveolar) and an increased proportion of smooth muscle cells in the lungs of CpcPH animals compared to controls and IpcPH (**Figure 1L**). To evaluate the potential effects of Mets on lung transcriptomic and control for potential biases related to the CpcPH model, we analyzed the transcriptomic profiles of Sham and Sham + Mets animals (**Figure S3A, B**). The comparative analysis revealed a modest number of DEGs, with only 27 DEGs detected; of these, four were upregulated and 23 were downregulated in the Sham + Mets group. This suggest that Mets has a relatively minimal impact on the lung transcriptome. Subsequently, we evaluated the impact of IpcPH and CpcPH on myocardial hypertrophy, fibrosis, and the recruitment of leukocytes and monocytes in both the left and right ventricles (**Figure S4A–H**). As anticipated, cardiomyocyte hypertrophy was evident in both the left and right ventricles in IpcPH and CpcPH models (**Figure S4A, E**). However, alongside the increased infiltration of inflammatory cells (monocytes and leukocytes), we observed that fibrosis and cardiomyocyte hypertrophy were more pronounced in both the RV and LV of CpcPH animals compared to IpcPH animals (**Figure S4A–H**).

Thus, our data support the existence of distinct pathophysiological phenotypes for IpcPH and CpcPH, at least in the preclinical setting. In CpcPH, the more severe pulmonary hemodynamic impairment appears to be driven by active pulmonary vascular remodeling and proliferative changes in pulmonary artery smooth muscle cells, as reflected by marked transcriptomic reprogramming and heightened lung inflammation. In contrast, elevated mPAP in IpcPH likely results predominantly from passive backward transmission of elevated left ventricular filling pressures, with minimal contribution from intrinsic pulmonary vascular alterations.

### Circulating Pro-inflammatory Factors Discriminate IpcPH from CpcPH Patients

In a translational effort, we investigated whether inflammation and circulating inflammatory markers could distinguish CpcPH from IpcPH in patients. We first analyzed complete blood counts and found no significant differences in white blood cell counts, neutrophils, lymphocytes, basophils, or the neutrophil-to-lymphocyte ratio between CpcPH and IpcPH patients (**Table 2**). We next profiled 96 circulating pro-inflammatory proteins using the Olink custom inflammation panel. Nine factors were differentially expressed between groups: eight were upregulated in CpcPH (TNF, IL-18, IL-12B, 4E-BP1, NT-3, NGF, FGF21, and FGF23), while CXCL5 was significantly downregulated (**Figure 2A–I**). To evaluate their diagnostic utility, we performed receiver operating characteristic (ROC) curve analysis, identifying HGF, TNF, and FGF23 as the most effective individual markers for distinguishing CpcPH from IpcPH, with area under the curve (AUC) values of 0.69, 0.69, and 0.68, respectively. To further improve discrimination, we conducted LASSO regression incorporating both readily available clinical variables (sex, diabetes status, dyslipidemia, age) and circulating biomarkers. This analysis identified a multivariate model, comprising diabetes status, FGF21, NT-3, CXCL5, HGF, and TNF, that achieved an AUC of 0.79 (**Figure 2J, S5A–C**). Among the nine proteins, IL-18 and 4E-BP1 were also upregulated at the transcriptomic level in the lungs of CpcPH animals compared to IpcPH (**Figure 2K, L and S5D-I**), and IL-18, 4E-BP1, FGF21, FGF23, HGF, IL12b, TNF, and NTF3 were elevated at the protein level in remodeled distal pulmonary vessels (**Figure 2M-U**). Notably, CXCL5 was not evaluated in the preclinical model due to the absence of a direct ortholog in rats.

**Figure 2.**
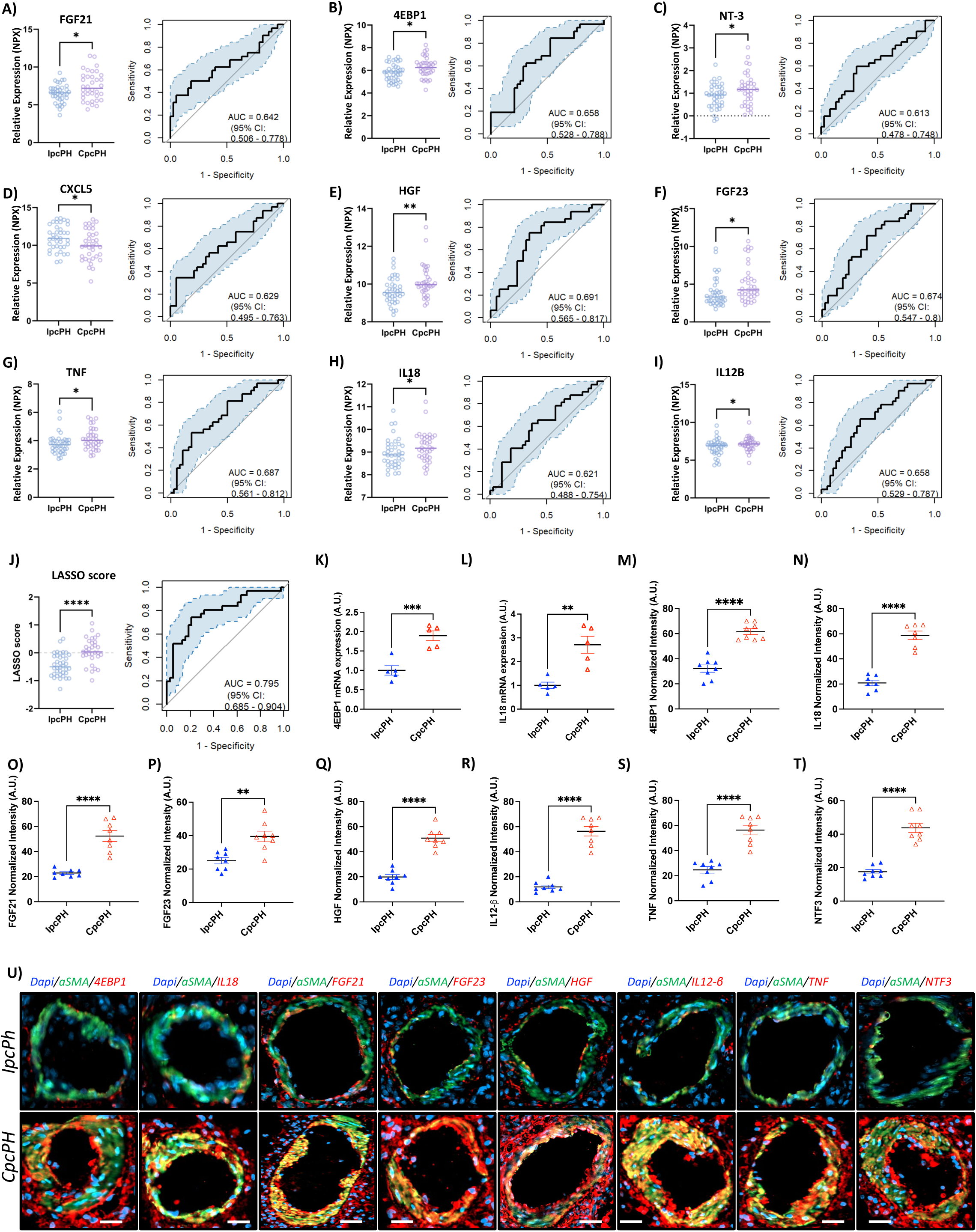
Inflammatory protein biomarkers distinguish IpcPH from CpcPH in Group 2 PH patients. (**A–I**) Relative plasma expression levels (left) and ROC curves (right) for nine inflammation-related proteins differentially expressed between CpcPH and IpcPH patients: **A**) FGF21, **B**) 4EBP1, **C**) NT-3, **D**) CXCL5, **E**) HGF, **F**) FGF23, **G**) TNF, **H**) IL-18, and **I**) IL-12B. Each protein showed modest-to-moderate discriminatory capacity (AUCs ranging from 0.642 to 0.691). **J**) LASSO-derived inflammation score significantly discriminates CpcPH from IpcPH and shows the highest AUC (0.795). **(K-L)** Transcriptomic expression levels of 8, 4E-BP1 in the lungs of IpcPH and CpcPH animals. **(M–T)** proteomic expression of IL-18, 4E-BP1, FGF21, FGF23, HGF, IL12b, TNF, and NTF3 in distal pulmonary arteries of IpcPH and CpcPH animals. Data are mean ± SEM; ns = not significant, *p < 0.05, **p < 0.01, ***p < 0.001, ****p < 0.0001. Representative immunofluorescence images illustrating their expression patterns are shown in **U)**. Scale bars = 50 µm

Then, we conducted a correlation analysis to assess the potential relationship between these proteins and disease severity parameters in human patients. Notably, IL-18, FGF23, HGF, 4E-BP1, FGF21, and the LASSO-score showed significant correlations with PVR, a key marker of pulmonary vascular remodeling (**Figure 3A**). Survival analysis revealed that NT-3, TNF, 4E-BP1, IL-12B, HGF, FGF21, FGF23, and the LASSO-score were significant predictors of mortality in univariate models and Kaplan-Meier analysis (**Figure 3B-I, Table S1**). To assess whether these proteins predict survival independently of the IpcPH/CpcPH subtype (given the distinct survival rates between these subgroups), we adjusted our analysis for IpcPH/CpcPH status. We found that FGF23, 4E-BP1, TNF, NT-3, and the LASSO-score predicted survival independently of subgroup status (model 1) (**Figure 3J, Table S2**). After further adjustment for sex, creatinine, and the 2022 ESC/ERS risk score (model 2), TNF and the LASSO-score remained the only independent predictors of survival (**Figure 3J, Table S2**). Finally, we assessed the incremental prognostic value of these inflammatory markers beyond NT-proBNP (**Figure S5 J-L**). NT-3, 4E-BP1, FGF23, and HGF provided additional prognostic value for survival.

**Figure 3.**
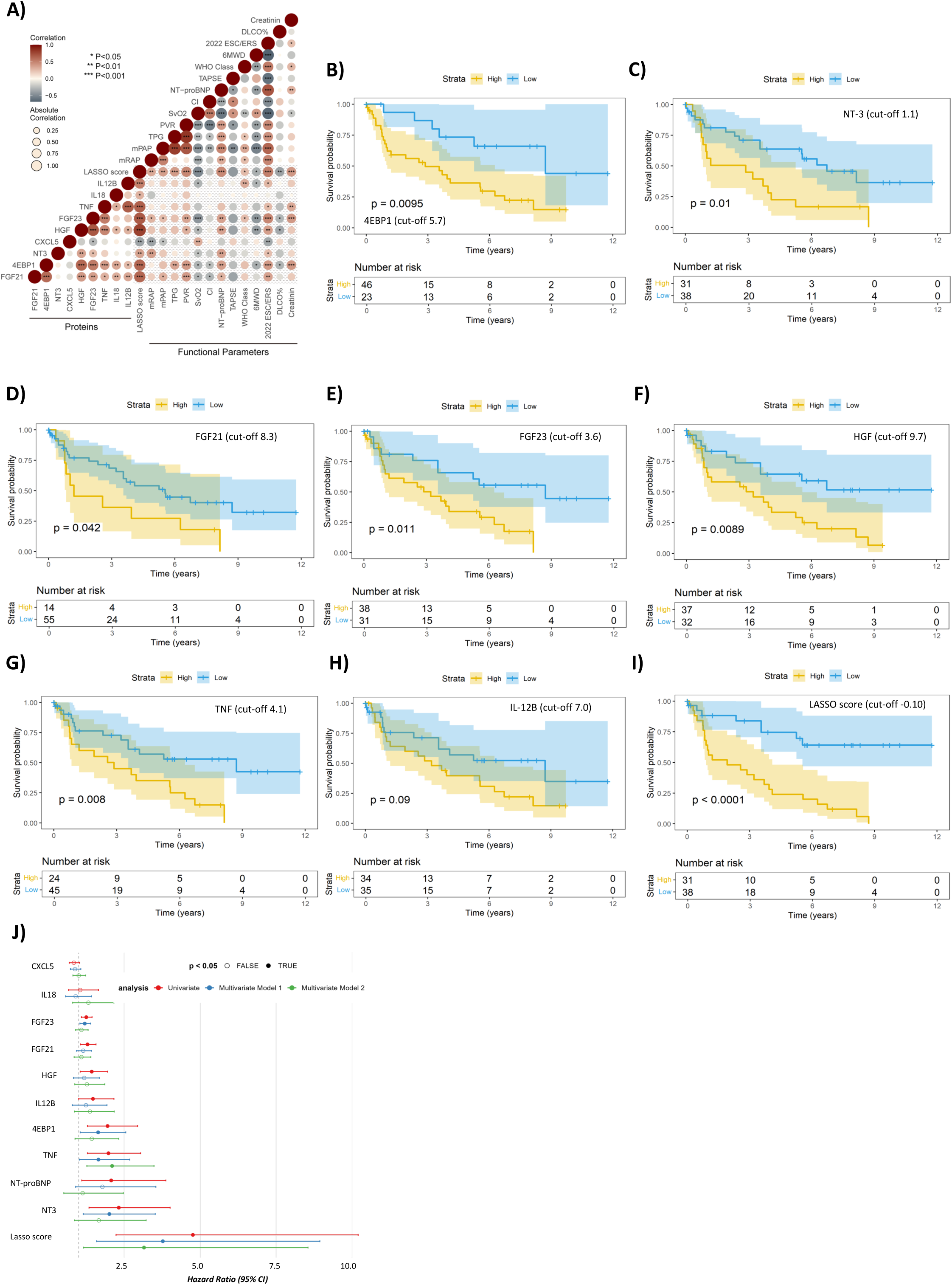
Prognostic relevance of inflammatory biomarkers in Group 2 PH. **A**) Correlation matrix between inflammatory markers, LASSO score, and clinical functional parameters. Circle size represents strength of correlation, color denotes direction, and significance levels are indicated. **B–I)** Kaplan–Meier survival curves stratified by high (yellow) vs low (blue) levels of individual biomarkers or composite scores, using optimal cut-offs: (B) 4E-BP1, (C) NT-3, (D) FGF21, (E) FGF23, (F) HGF, (G) TNF, (H) IL-12B, and (I) LASSO-derived inflammatory score. **J**) Forest plot displaying hazard ratios (HRs) and 95% confidence intervals for each inflammatory marker in three Cox regression models: univariate (red), multivariate model 1 (blue; adjusted for IpcPH/CpcPH status), and multivariate model 2 (green; additionally adjusted for sex, creatinine, and the ESC/ERS 2022 risk score). Filled circles indicate markers significantly associated with survival (p < 0.05), while open circles denote non-significant associations. All statistical comparisons performed using Mann–Whitney or unpaired t tests as appropriate. AUC: area under the curve; ROC: receiver operating characteristic; A.U.: arbitrary units.

### Levosimendan improves pulmonary haemodynamic parameters in IpcPH and CpcPH preclinical model

We subsequently evaluated the impact of levosimendan administration (3mg/kg/day for 3 weeks) on animals with IPCPH and CpcPH. As anticipated, levosimendan notably enhanced LV systolic function, as evidenced by increased MAPSE and LV-S wave, alongside improvements in global cardiac function, including increased CO and SV (**Table 1**). It is noteworthy that due to the fixed pressure elevation induced by banding, we did not observe significant changes in TBG or LVSP. However, a trend toward improved LV diastolic function was observed, indicated by decreased LVEDP and E/E’ ratio in both IpcPH and CpcPH. Furthermore, levosimendan administration led to enhancements in RV function, as evidenced by decreased RVEDP, increased TAPSE, RV-S wave, and improved TAPSE/RVSP coupling ratio. Additionally, levosimendan demonstrated favorable effects on pulmonary hemodynamics, characterized by increased PAAT and decreased RVSP, PVR, and TPG. The longitudinal echocardiographic follow-up indicates that the impact of levosimendan on hemodynamics becomes evident within one week of treatment initiation in both IpcPH and CpcPH animal models (**Figure S6A-N**). Additionally, we observed no significant differences in weight between treated and non-treated animals, suggesting good tolerance to the treatment (**Figure S6G, N**). Our subsequent investigations focused on evaluating the effects of levosimendan on the histology of the LV, RV, and pulmonary tissues. Consistent with the improvements observed in the hemodynamic parameters of the LV and RV, treatment with levosimendan resulted in a notable reduction in fibrosis, cardiomyocyte hypertrophy, and inflammatory cell infiltration, mainly in CpcPH (**Figure S7A-H**). Additionally, levosimendan ameliorated adverse remodeling of the pulmonary vasculature, reduced PASMC proliferation, and promoted PASMC apoptosis (**Figure 4A-D**). These effects were associated with a significant decrease in monocyte and leukocytes infiltration, specifically in the lungs of CpcPH animals (**Figure 4A, E, F**). However, in IpcPH models, levosimendan did not show any discernible impact on pulmonary vasculature or the structures of the RV and LV.

**Figure 4.**
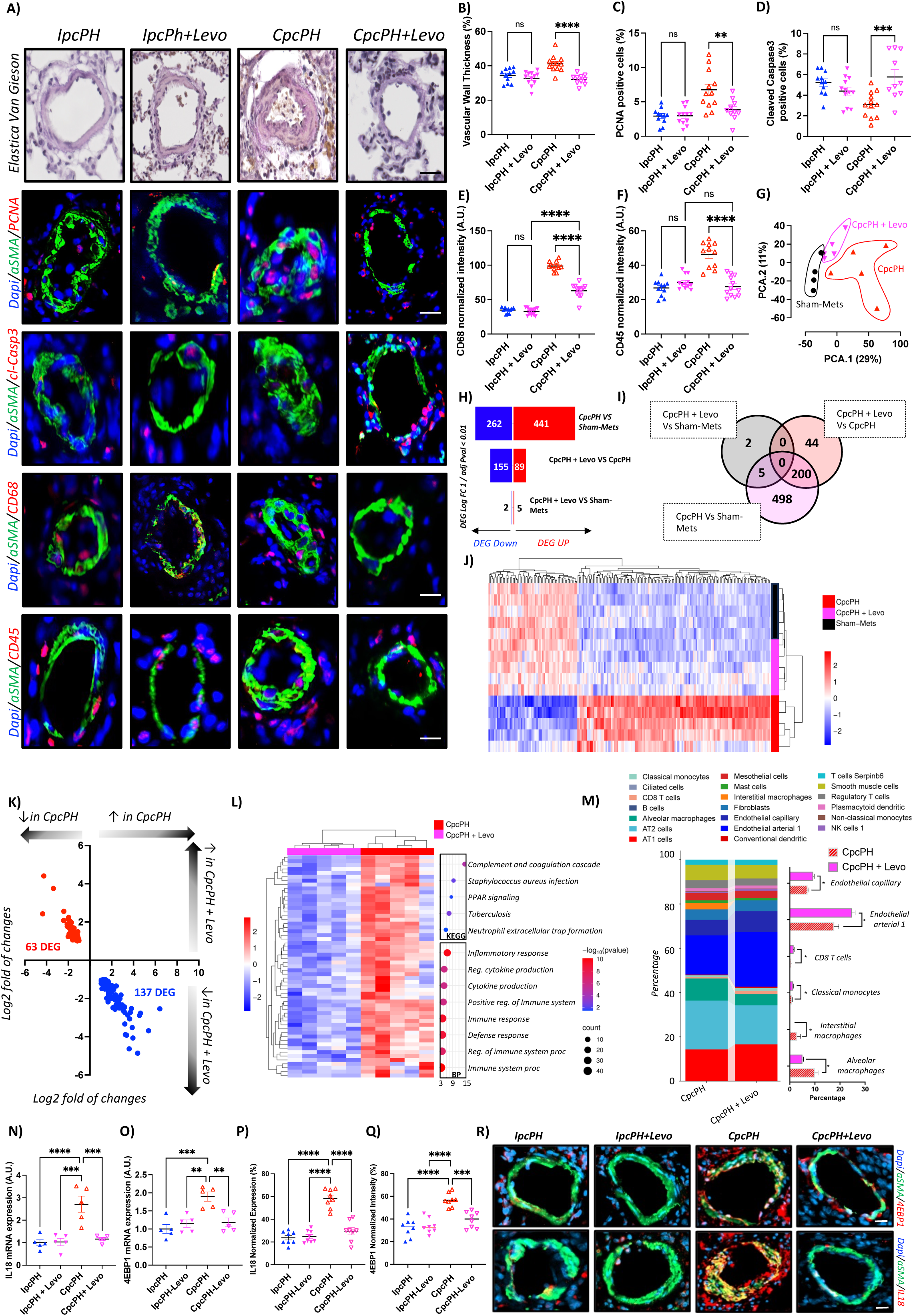
Levosimendan attenuates pulmonary vascular remodeling, inflammation, and transcriptomic reprogramming in a preclinical model of CpcPH. **A**) Representative histological and immunofluorescence images of lung sections from IpcPH, IpcPH + Levosimendan, CpcPH, and CpcPH + Levosimendan animals. Elastica Van Gieson staining shows medial thickening in CpcPH lungs, which is attenuated by Levosimendan. Immunofluorescence images illustrate PASMC proliferation (α-SMA⁺/PCNA⁺), apoptosis (α-SMA⁺/Cleaved Caspase-3⁺), monocyte (CD68⁺), and leukocyte (CD45⁺) infiltration. Scale bars = 20 **µm.** Quantification of: **B**) pulmonary vascular medial thickness, **C**) proliferating PASMCs (PCNA⁺), **D**) apoptotic PASMCs (Cleaved Caspase-3⁺), **E**) CD68⁺ and **F**) CD45⁺ fluorescence intensity. n = 5–8/group. Data are shown as mean ± SEM. ns = not significant, ***p < 0.001, ****p < 0.0001. **G**) Principal component analysis (PCA) of lung transcriptomes showing distinct clustering of Sham-Mets, CpcPH, and CpcPH + Levosimendan. **H**) Differentially expressed genes (DEGs; log2FC > 1, adj. p < 0.05) across conditions. Red and blue bars represent upregulated and downregulated DEGs, respectively. **I**) Venn diagram showing the overlap of DEGs reversed by Levosimendan in CpcPH animals. **J)** Heatmap with unsupervised clustering of the 200 DEGs identified in CpcPH animals compared to their respective Sham controls that are also modulated by levosimendan treatment, illustrating partial normalization of the CpcPH transcriptomic signature following therapy. **K**) Quadrant plot depicting genes differentially expressed in CpcPH lungs and reversed by levosimendan treatment. The X-axis represents log2 fold changes in CpcPH versus Sham + Mets, and the Y-axis represents log2 fold changes in CpcPH + Levosimendan versus untreated CpcPH. A total of 137 genes were downregulated and 63 upregulated by levosimendan, indicating partial reversal of the CpcPH transcriptomic profile. **L**) KEGG and GO enrichment analysis of reversed DEGs demonstrates suppression of inflammation-related pathways (e.g., complement cascade, cytokine signaling, innate immune response) by Levosimendan. **M**) CIBERSORTx-based cell deconvolution shows changes in lung immune and endothelial cell populations in CpcPH and after Levosimendan treatment, including reductions in alveolar and interstitial macrophages and increases in endothelial cells and CD8⁺ T cells. **N, O)** Transcriptomic and **P, Q)** proteomic expression levels of IL-18 and 4EBP1 in the lungs of IpcPH and CpcPH animals, with and without levosimendan treatment. **R)** Representative immunofluorescence images, and related quantification showing IL-18 and 4EBP1 expression in pulmonary arteries, highlighting increased expression in CpcPH animals and partial reversal following levosimendan therapy. Scale bars = 50 µm. All statistical comparisons performed using Mann–Whitney or unpaired t tests as appropriate. AUC: area under the curve; ROC: receiver operating characteristic; A.U.: arbitrary units.

### Levosimendan Reverses Adverse Lung Transcriptomic Reprogramming Observed in a Preclinical Model of CpcPH

To further understand the molecular mechanisms underlying the effects of levosimendan, we undertook a transcriptomic analysis of lung tissues from both CpcPH and IpcPH models treated with levosimendan. PCA revealed no distinct clustering between control (Sham), IpcPH, and IpcPH animals treated with levosimendan (**Figure S8A**), suggesting a lack of significant transcriptomic changes between those conditions. Additionally, our analysis did not identify any DEGs (log FC>1 and FDR< 0.05) when comparing Sham to IpcPH, as well as IpcPH to IpcPH animals treated with levosimendan. This lack of significant DEGs suggests that, from a transcriptomic perspective, levosimendan does not induce notable molecular alterations in the lungs of IpcPH models. In contrast, the PCA revealed distinct clustering of the transcriptomes for the control (Sham + Mets), CpcPH, and CpcPH treated with levosimendan groups, clearly delineating them according to their respective conditions in an unsupervised manner (**Figure 4G**). We observed 703 DEGs when comparing CpcPH to sham + Mets with the 441 upregulated DEG being mainly related to inflamation and the 262 dowregulated DEGS being related to Ion channel activity based on GO analysis (**Figure 4H and S9A-C**). Interestingly the 155 DEGs downregulated in the lungs of CpcPH animals treated with levosimendan, as compared to untreated CpcPH animals, are predominantly associated with inflammatory processes while the upregulated DEGs appear to be involved in fatty acid metabolism (**Figure 4H and Figures S10 A-C**). Surprisingly, only 7 DEGs were observed when comparing levosimendan-treated CpcPH animals to the control (Sham + Mets) group, (**Figure 4H and Figures S11 A-B**). Then, we evaluated the overlap of DEGs across all conditions to identify transcripts in the lungs affected by CpcPH that are also influenced by Levosimendan treatment (**Figure 4I**). Our analysis revealed that 200 transcripts altered in CpcPH lungs compared to controls are also affected in Levosimendan-treated CpcPH lungs. Notably, we found that 137 DEGs upregulated in CpcPH are downregulated with treatment, and 63 DEGs downregulated in CpcPH are upregulated by Levosimendan, suggesting a partial reversal of CpcPH-associated transcriptomic changes by the treatment (**Figure 4J, K)**. GO analysis of these DEGs indicates that the biological functions primarily associated with the 137 downregulated DEGs are related to inflammation, whereas the upregulated DEGs are linked to fatty acid metabolism (**Figure 4L, S12A-C)**. To identify potential upstream regulators of the transcriptomic changes induced by levosimendan, we performed transcription factor enrichment analysis on the 137 genes that were upregulated in CpcPH and downregulated following treatment. This analysis revealed NF-κB as the only significantly enriched transcription factor (**Figure S13A**). In line with this finding, we observed reduced NF-κB activation in the lungs of levosimendan-treated CpcPH animals (**Figure S13B**). Consistent with our histological observations (**Figure 4A, E, F**), cellular deconvolution analysis of lung transcriptomes revealed that levosimendan treatment was associated with a reduction in macrophage infiltration (both interstitial and alveolar), along with increased representation of classical monocytes, capillary and arterial endothelial cells, and CD8⁺ T cells compared to untreated CpcPH animals (**Figure 4M**). Next, we evaluated the lung expression of the nine circulating inflammatory markers that were differentially expressed between CpcPH and IpcPH patients. Among these, only IL-18 and 4E-BP1 were significantly upregulated at the transcriptomic level in the lungs of CpcPH animals and were reduced following levosimendan treatment (**Figure 4N, O**). At the protein level, both IL-18 and 4E-BP1 as well as FGF21, FGF23, HGF, IL12b, TNF, and NTF3 were elevated in distal pulmonary arteries of CpcPH animals and decreased in parallel with the attenuation of pulmonary vascular remodeling and inflammation after levosimendan treatment (**Figure 4P–R and S14A-G**). These findings collectively suggest that levosimendan not only mitigates the adverse pulmonary vascular remodeling observed in CpcPH lungs but may also exert its effects by reprogramming maladaptive transcriptomic changes and reducing inflammation.

## Discussion

In this multicentric preclinical study, we first validated two distinct animal models of Group-2PH, reflecting the human hemodynamic definitions of IpcPH and CpcPH. As expected, CpcPH animals exhibited more severe pulmonary vascular remodeling, increased inflammatory cell infiltration, and marked transcriptomic alterations compared to IpcPH animals. To translate these findings to patients, we identified nine circulating inflammation-related proteins and cytokines that successfully distinguished CpcPH from IpcPH, suggesting potential use as biomarkers for disease subtyping. We next demonstrated that levosimendan treatment improved right and left ventricular function and favorably impacted pulmonary hemodynamics in both models. However, only in CpcPH animals did levosimendan significantly reduce pulmonary vascular remodeling and lung inflammation, alongside a reversal of the disease-associated transcriptomic reprogramming. Transcription factor analysis revealed NF-κB as the top enriched upstream regulator of the CpcPH gene signature, and its activation was decreased following levosimendan treatment, pointing to a mechanistic role in its anti-inflammatory effects. In contrast, levosimendan had minimal molecular or histological impact on IpcPH lungs. Among the nine inflammatory markers, IL-18 and 4E-BP1 stood out for being elevated in both the plasma and lungs of CpcPH animals, and their expression was reduced after levosimendan treatment. Importantly, 4E-BP1 also independently predicted survival in patients with IpcPH and CpcPH (**Figure 5**).

**Figure 5.**
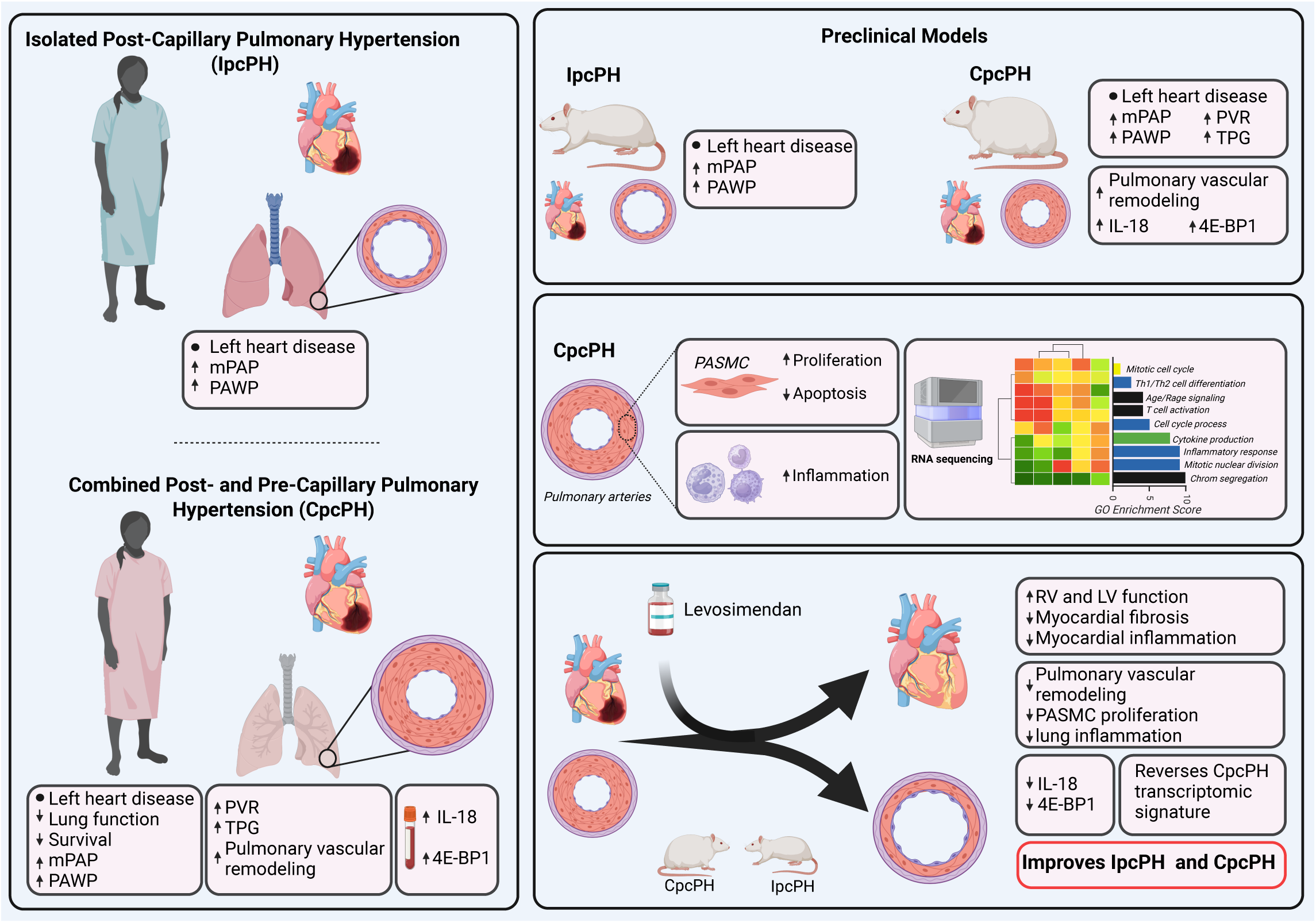
Graphical Abstract. Group 2 pulmonary hypertension (PH) includes isolated post-capillary PH (IpcPH) and combined pre-and post-capillary PH (CpcPH), which differ in pulmonary vascular involvement and clinical outcomes. In a well-characterized patient cohort, CpcPH was associated with worse lung function, more severe hemodynamics, elevated circulating IL-18 and 4E-BP1, and reduced survival compared to IpcPH. These clinical phenotypes were faithfully recapitulated in translational rodent models. CpcPH animals displayed marked pulmonary vascular remodeling, enhanced PASMC proliferation, reduced apoptosis, inflammation, and a distinct transcriptomic profile. Levosimendan treatment improved biventricular function and reduced myocardial fibrosis, pulmonary vascular remodeling, and inflammation. It also normalized IL-18 and 4E-BP1 expression and reversed the maladaptive transcriptomic signature in CpcPH lungs. These findings suggest that levosimendan exerts dual therapeutic effects, on both cardiac performance and pulmonary vascular pathology, supporting its relevance as a potential disease-modifying treatment in both IpcPH and CpcPH.

It is intriguing to note that while levosimendan improves pulmonary hemodynamics in both IpcPH and CpcPH models, it exerts molecular and histological effects exclusively in the lungs of CpcPH animals. Our data, coupled with insights from existing literature, suggest that levosimendan ameliorates PH through two distinct mechanisms. Given its well-established role as a calcium sensitizer used in the management of heart failure, particularly for enhancing myocardial contractility, one mechanism likely involves the improvement of left and right ventricular function. This is supported by evidence from patients with left heart failure as well as preclinical models of right heart failure induced by PAH or pressure overload (pulmonary artery banding) (11–14,16,17). In these contexts, levosimendan has been shown to enhance both left and right ventricular systolic and diastolic function, reduce myocardial fibrosis, and increase cardiac output. Additionally, as a calcium sensitizer, levosimendan may exert direct effects on pulmonary vascular tone. It is proposed to reduce calcium sensitivity of contractile proteins in vascular smooth muscle, lower intracellular free calcium levels, potentially inhibit phosphodiesterase III, and open potassium channels, collectively promoting vasodilation and vascular relaxation(26). These favorable effects on the cardiovascular system may underlie the therapeutic efficacy of levosimendan in treating IpcPH. Conversely, our findings indicate that levosimendan exerts a beneficial effect on adverse pulmonary vascular remodeling by reducing vascular wall thickness, PASMC proliferation, and promoting PASMC apoptosis. This observation resonates with previous research conducted in preclinical models of Group-1 PH, where three weeks of levosimendan treatment mitigated the development of PAH by inhibiting PASMC proliferation and reducing pulmonary vascular medial wall thickness(16,17).

At the molecular level, our transcriptomic analysis reveals that levosimendan is associated with decreased expression of genes implicated in inflammatory processes as well as decreased accumulation of leukocytes and monocytes. This aligns with findings from Revermann and colleagues, who observed an anti-inflammatory effect of levosimendan on the lung vasculature(16). Beyond the lung, a metanalysis focusing on the reported anti-inflammatory effect of levosimendan in clinical trial in advanced heart failure reported that levosimendan is associated with decreased expression of pro-inflammatory cytokine IL-6 and mPAP(27). Leveraging from literature in the group 1PH, IL-6 is a key driver of adverse pulmonary vascular remodeling in PAH. Interestingly, in our dataset, IL-6 is increased in the lung of CpcPH animals and decreased following levosimendan, at least from the transcriptomic standpoint. Moreover, the literature suggests that Levosimendan decreases IL-1β and induces increased IL-6 expression by blocking NF-KB activation(28). Consistent with the anti-inflammatory effect of the levosimendan, levosimendan improves survival in the preclinical setting of sepsis(29). Although controversial in the literature, in patients with severe sepsis and septic shock or sepsis-induced myocardial dysfunction, levosimendan is associated with a significant reduction in mortality compared with standard inotropic therapy(30,31). Thus, beyond the inotropic effect of the levosimendan, an anti-inflammatory effect might contribute to the therapeutic effect of levosimendan At the molecular level, our transcriptomic analysis demonstrates that levosimendan treatment is associated with decreased expression of genes involved in inflammatory pathways, along with reduced accumulation of leukocytes and monocytes in the lungs. These findings align with those of Revermann et al., who reported an anti-inflammatory effect of levosimendan on pulmonary vasculature in preclinical models(16). Beyond the pulmonary context, a meta-analysis of clinical trials in patients with advanced heart failure revealed that levosimendan significantly reduced circulating levels of the pro-inflammatory cytokine IL-6, as well as mPAP (27). Drawing from the literature in Group 1 PH, IL-6 is recognized as a key driver of adverse pulmonary vascular remodeling in PAH(32). Consistent with this, we observed elevated IL-6 expression in the lungs of CpcPH animals, which was reduced following levosimendan treatment, at least at the transcriptomic level. Moreover, the literature suggests that levosimendan may exert its anti-inflammatory effects, in part, by inhibiting NF-κB activation and consequently reducing IL-1β-induced IL-6 production(28). Supporting this anti-inflammatory profile, levosimendan improves survival in preclinical models of sepsis(29). Although somewhat controversial, clinical studies in patients with severe sepsis, septic shock, or sepsis-induced myocardial dysfunction suggest that levosimendan is associated with reduced mortality compared with standard inotropic therapies (30,31). Altogether, these data raise the possibility that, beyond its well-established inotropic effects, the anti-inflammatory properties of levosimendan may significantly contribute to its therapeutic efficacy in pulmonary hypertension. Conflicting literature suggests that levosimendan may exert vasodilatory effects on pulmonary arteries, potentially through the opening of potassium channels, as shown in ex vivo experiment(18). Additionally, systemic veno dilation has been reported upon levosimendan treatment(9). However, in vivo studies in preclinical models of Group-1 PH, and our own transcriptomic analyses in IpcPH and CpcPH, did not reveal significant modulation of ion channel expression or direct evidence of altered pulmonary vasoreactivity. Overall, our results suggest that levosimendan exerts its therapeutic effects in PH through a dual mechanism involving improved biventricular cardiac function and attenuation of adverse pulmonary vascular remodeling, thereby offering potential avenues for targeted therapy in different forms of the disease.

Both IpcPH and CpcPH are characterized by elevated mPAP and PAWP. In most patients with IpcPH, elevated mPAP can be attributed to increased LV filling pressure, although it may also be influenced by pulmonary vascular remodeling. Similarly, preclinical models of IpcPH demonstrate moderate pulmonary vascular remodeling without significant increases in PASMC proliferation, decreases in PASMC apoptosis, or substantial molecular changes in lung tissue. Although intriguing, this may reflect the passive nature of vascular remodeling, involving subtle changes that may go undetected in whole lung transcriptome analysis. Conversely, pulmonary artery remodeling is a prominent feature in CpcPH, resembling the pattern observed in PAH. In preclinical models of CpcPH, substantial pulmonary vascular remodeling is evident, characterized by increased PASMC proliferation, decreased PASMC apoptosis, and increased inflammation. Transcriptomic analysis reveals a pro-inflammatory state in whole lung tissue, along with decreased expression of ion channels, suggesting impaired vasoreactivity in CpcPH lungs. Furthermore, comparison of the transcriptomic signatures between CpcPH and IpcPH lungs indicates that, similar to PAH, upregulated DEGs in CpcPH are associated with cell proliferation and the cell cycle, while downregulated DEGs are associated with lymphocyte activation and function. Interestingly, this observation echoes recent findings that impaired T cell signaling contributes to adverse pulmonary vascular remodeling in preclinical models of Group-2 PH, regardless of whether the model is of IpcPH or CpcPH(33,34). These findings suggest that IpcPH and CpcPH arise from distinct etiologies, with CpcPH sharing similarities with the molecular dysfunction observed in PAH. However, echoing clinical trials that do not differentiate between CpcPH and IpcPH, and irrespective of the underlying disease etiology (whether involving active vascular remodeling or not), Levosimendan alleviates PH in both CpcPH and IpcPH animal models.

Based on our preclinical observations, inflammation appears to be a key determinant of CpcPH, and, and, consistent with literature on Group 1 PH, a critical driver of adverse pulmonary vascular remodeling(35). Although we did not observe substantial differences in circulating inflammatory cell counts between IpcPH and CpcPH patients, we identified nine circulating inflammation-related proteins that were differentially expressed in CpcPH compared to IpcPH patients. Among them, IL-18 has previously been reported to be elevated in the blood of Group-2 PH patients, although without distinguishing IpcPH from CpcPH(33,34). Mechanistically, these studies demonstrated that IL-18 accumulation originates from altered lymphocyte function and that blocking IL-18 ameliorates pulmonary vascular remodeling in preclinical models of Group-2 PH. Similarly, in our study, we observed that levosimendan treatment, in parallel with reduced pulmonary vascular remodeling, also decreased IL-18 expression. Drawing from the Group-1 PH literature, many of the nine circulating proteins, such as FGF21, 4EBP1, HGF, and TNF, have been linked to pulmonary vascular remodeling(36–39). Additionally, FGF23 has been associated with right ventricular dysfunction and PAH(40), while CXCL5 has been implicated in systemic vascular remodeling(41). Notably, FGF21, NT-3, 4EBP1, CXCL5, HGF, TNF, IL-18, and IL-12b have also been associated with heart failure or cardiovascular disease(42–50). Whether, in human patients, the differential expression of these inflammation-related proteins in CpcPH reflects worse RV/LV function and adverse pulmonary vascular remodeling and hemodynamics remains to be determined. However, given their increased expression in the lungs of CpcPH animals, alongside decreased expression following levosimendan treatment and corresponding improvements in vascular remodeling, we speculate that IL-18 and 4E-BP1 dysregulation in CpcPH patients may mirror adverse pulmonary vascular lesions.

In conclusion, our study characterized two preclinical models of Group-2PH that faithfully replicate the human IpcPH and CpcPH phenotypes from a hemodynamic perspective. Through this investigation, we uncovered distinct pathophysiological and molecular signatures distinguishing IpcPH from CpcPH. IpcPH primarily arises from passive elevation of left ventricular filling pressures and exerts only a moderate impact on lung vasculature. In contrast, CpcPH is marked by severe pulmonary vascular remodeling, significant inflammatory cell infiltration, and a unique inflammatory lung transcriptomic profile. Our study extends beyond hemodynamic definitions, underscoring how the differences between IpcPH and CpcPH, including the heightened inflammation in CpcPH, profoundly shape lung vascular pathology in the preclinical setting. The primary aim of our work was to assess the effects of levosimendan on lung vasculature in Group-2 PH. Beyond its well-known benefits on left and right ventricular function, we found that levosimendan significantly attenuated adverse pulmonary vascular remodeling in the lungs of CpcPH animals. This beneficial effect is likely mediated by levosimendan’s anti-inflammatory properties. In a translational effort, we identified nine circulating inflammation-related proteins differentially expressed between IpcPH and CpcPH patients. Among them, IL-18 and 4E-BP1 were upregulated in the lungs of CpcPH animals and decreased following levosimendan treatment in animal models. Notably, 4E-BP1 was also predictive of survival in human patients. From a molecular standpoint, our findings suggest that inflammation may play a critical role in the development of pulmonary vascular lesions in CpcPH, and circulating inflammatory proteins may have potential as biomarkers to discriminate IpcPH from CpcPH in patients. Finally, our work offers fresh insights into the promising therapeutic effects of levosimendan observed in ongoing clinical trials for Group-2 PH. We propose that levosimendan holds significant potential to improve outcomes in both IpcPH and CpcPH by enhancing cardiac function and mitigating adverse pulmonary vascular remodeling.

### Limitation

Our study encountered several limitations that merit consideration. Firstly, the unavailability of human lung samples from patients with IpcPH and CpcPH restricted our ability to directly assess transcriptomic and histological differences in these populations. Compounding this issue is the challenge of obtaining post-mortem lung samples, further exacerbated by the occurrence of flash pulmonary edema in patients with terminal left heart failure, potentially compromising tissue integrity and molecular profiles. Additionally, in our animal model, distinguishing remodeled small arteries from remodeled small veins posed a significant challenge, despite utilizing elastica van Gieson staining for discrimination. This limitation underscores the complexity of accurately interpreting histological findings in the absence of well-defined arterial and venous specific markers. Furthermore, while the literature did not demonstrate significant effects of Levosimendan on metabolic syndrome parameters, including body weight and glucose tolerance, we cannot rule out the possibility that the therapeutic effects of Levosimendan on lung vasculature may be influenced by improvements in metabolic syndrome.

