## supplemental methods and figures for "Levosimendan Ameliorates Adverse Pulmonary Vascular Remodeling in Group-2 Pulmonary Hypertension"

**Supplemental Methods and legend:**

***Histology and biochemistry***

Paraffin-embedded lung sections from rats (5 µm) were labelled with antibodies against α-SMA to measure PA medial thickness (1:400, A2547, Sigma-Aldrich, Oakville, ON, Canada; 1:400, ab5694, Abcam, Toronto, ON, Canada), Ki67 for proliferation (1:200, AB9260, MilliporeSigma, Bilerica, MA, USA), and anti-cleaved caspase-3 for apoptosis (1:150,#9661, cell signaling, Danvers, USA). Additional primary antibodies were used to characterize immune and molecular markers in our samples: anti-CD45 (1:200, Cell Signaling Technology, Cat# 13917, Danvers, USA) to identify total leukocytes, anti-CD68 (1:200, Cell Signaling Technology, Cat# 76437, Danvers, USA) to detect macrophages, anti-4EBP1 (1:200, #9644, cell signaling, Danvers, USA) to detect the eukaryotic translation initiation factor 4E-binding protein 1 and anti-IL-18 (1:200, ab191860, Abcam, Toronto, ON, Canada) to evaluate pro-inflammatory cytokine levels. For immunofluorescence, antibody binding was detected with secondary antibodies at 1:1000 in PBS: goat anti-rabbit Alexa Fluor® 488 (1:1000, Life Technologies A-11008), goat anti-mouse Alexa Fluor 594 (1:1000, Life Technologies A-11005), goat anti-rabbit Alexa Fluor® 594 (1:1000, Life Technologies A-11037), and goat anti-mouse Alexa Fluor® 488 (1:1000, Life Technologies A-11001). The nucleus was labeled with DAPI (4′,6-diamidino-2-phenylindol) Fluoromount G mounting medium (Life Technologies, Burlington, CAN). Image acquisition was performed on a Zeiss Apotome.2 imaging microscopy workstation (Zeiss, Oberkochen, Germany) using multiple fluorescence channels. Zen Lite software (Zeiss, Oberkochen, Germany) was used to semi-automatically quantify the images. 15 intra-acinar pulmonary arteries/rat were examined. Indices were calculated as the percentage of positive-staining cells among DAPI-stained nuclei/artery. The mean of these measurements was used as the representative value for each animal. For immunohistochemistry, Elastica Van Gieson staining (Elastin solution, HX98471391, Millipore; Hematoxylin alcohol and iron chloride solutions; Picrofushin solution, HX99264299, Millipore) was conducted according to the supplier's recommendations to assess pulmonary vascular remodeling. Fifteen random arteries (diameter < 50 µm) per rat were investigated. The percentage of remodeling was assessed by the ratio of artery external to its internal diameter. Elastic fibers were visualized using Elastin stain, which colors the fibers black/purple; Weigert's hematoxylin stain was used to visualize the nuclei in brown/black. Paraffin-embedded RVs were sectioned at 5µm and stained with hematoxylin-eosin (for RV hypertrophy) and Masson-trichrome (for fibrosis). Cardiomyocyte cross-sectional area was determined by tracing the outlines of cardiomyocytes with a clear nucleus image in hematoxylin-eosin-stained sections, as previously described. Quantification of RV fibrosis and cross-sectional area (CSA) of cardiomyocytes was performed in at least 10 randomly chosen areas using ImageJ software.

***Echocardiography***

Non-invasive longitudinal assessment of pulmonary, right ventricular, and left ventricular function was conducted post-surgery until the end of the protocol. Transthoracic echocardiography was performed using a Vevo® 2100 (FUJIFILM VisualSonics Inc.) phased-array, color-Doppler ultrasound system equipped with a 37.5 MHz transducer, capable of frame rates up to 1000/s. Serial 2-dimensional, M-mode, and pulsed-wave Doppler ultrasound recordings were obtained under anesthesia (inhaled isoflurane, 1.6–2.0%, mixed with humidified medical air delivered via a cone inhaler). Mitral annular plane systolic excursion (MAPSE), left ventricular (LV) S-wave, and E/E' ratio served as indicators of LV systolic and diastolic function. LV ejection fraction (LVEF) was calculated using the modified Quinones equation. Tricuspid annular plane systolic excursion (TAPSE), right ventricular (RV) S-wave, and pulmonary artery acceleration time (PAAT) were quantified to assess RV systolic and diastolic function and PH, as described in(21).

***Right and Left Heart Catheterization.***

Invasive, closed-chest, right heart and left heart catheterization were performed using the Transonic Scisense ADV500 Pressure-Volume Measurement System and analyzed with Labscibe2 software from Work System, Inc, following previously described methods(21,22). Measurements included RV and LV systolic pressures (RVSP; LVSP), LV end-diastolic pressure (LVEDP), stroke volume (SV), and cardiac output (CO). Mean pulmonary arterial pressure (mPAP) was estimated using the Chemla equation (0.61*RVSP + 2)(23). PVR were determined by the equation PVR=(mPAP-LVEDP)/CO, and TPG was calculated as TPG=mPAP-LVEDP. RV-pulmonary artery coupling was estimated by the TAPSE/RVSP ratio(24). Constriction severity was estimated by the difference of trans-stenosis pressures gradient (TPG = pre-banding pressure-post-banding pressure).

**RNA-sequencing**

Alignment of FASTQ files to the mRatBN7.2 rat reference genome, along with gene annotations from Ensembl build 98, was performed using STAR (v2.6.1a)(25). Gene quantification was conducted using RSEM (v1.3.1). Quality control analysis involved multiQC. Downstream analysis was carried out using R 4.0. Each tissue was analyzed separately, with initial gene filtering performed using the filterByExpr() function from edgeR. Subsequent differential expression analysis utilized the DESeq2 protocol(26). Unsupervised sample clustering was achieved through principal component analysis (PCA).

**Figure S1. Experimental design of preclinical models of Group 2 pulmonary hypertension (PH-LHD). A**) Experimental timeline for the IpcPH model (Model-1) developed at Queen's University. Male Sprague Dawley rats underwent sham or supracoronary aortic banding (SAB) surgery at Week 0. Levosimendan (3 mg/kg/day, oral gavage) or vehicle was administered from Week 7 to Week 10. Serial echocardiography (Echo) was performed at Weeks 7, 9, and 10. Right heart catheterization (RHC) and tissue collection were performed at Week 10. **B**) Experimental timeline for the CpcPH model (Model-2) established at PHRG-Québec. Rats underwent SAB at Week 0 and were fed a high-fat diet combined with olanzapine (4 mg/kg IP every 2 days) from Week 1 to Week 10. Animals received vehicle or levosimendan (3 mg/kg/day, oral gavage) from Week 7 to Week 10. Echo was performed at Weeks 7, 9, and 10. RHC and tissue collection occurred at Week 10.

**Figure S2. Survival analysis of patients with IpcPH and CpcPH. A**) Kaplan–Meier survival curve comparing transplant-free survival in patients diagnosed with isolated post-capillary pulmonary hypertension (IpcPH, blue) and combined pre- and post-capillary pulmonary hypertension (CpcPH, red). The analysis demonstrates significantly worse survival in CpcPH patients compared to IpcPH patients (log-rank P = 0.0146; hazard ratio [HR] = 0.4575; 95% confidence interval: 0.2443–0.8568). The shaded areas represent the 95% confidence intervals. The number of patients at risk at each time point is shown below the x-axis.

**Figure S3. Transcriptomic impact of metabolic syndrome (MetS) in control animals.** **A)** Volcano plot showing differentially expressed genes (DEGs) in lungs from Sham-Mets rats compared to Sham controls. Genes significantly upregulated are shown in red, and those downregulated are shown in green (adjusted $p < 0.01$ and $|\log_2 \text{fold change}| > 1$). **B)** Heatmap of the top DEGs between Sham-Mets and Sham animals, clustered by gene expression Z-scores. Data suggest a relatively modest impact of MetS alone on lung transcriptomic profiles.

**Figure S4. CpcPH is associated with increased cardiomyocyte hypertrophy, fibrosis, and inflammatory cell infiltration in both the left and right ventricles. (A–D) Left ventricle (LV):** **A)** Quantification of cardiomyocyte cross-sectional area with representative hematoxylin and eosin (H&E) staining. **B)** Quantification of LV fibrosis with representative Masson's trichrome staining. **C)** Representative immunofluorescence images and quantification of $CD68^+$ monocytes/macrophages and **D**) $CD45^+$ leukocytes in the LV myocardium. **(E–H) Right ventricle (RV): E)** Cardiomyocyte cross-sectional area and H&E staining. **F)** Quantification of RV fibrosis and corresponding Masson's trichrome staining. **G)** Immunofluorescence and quantification of $CD68^+$ **H)** and $CD45^+$ immune cell infiltration in the RV. Data are presented as mean ± SEM. Statistical significance was determined by one-way ANOVA followed by Tukey's multiple comparison test. *P < 0.05, **P < 0.01, ****_P < 0.0001, ns = not significant._

**Figure S5. Prognostic modeling of inflammatory proteins and LASSO-derived score in CpcPH patients**. **A)** Table of variables included in the multivariable LASSO regression model for diagnosis prediction. **B)** Cross-validation plot showing the optimal λ value (vertical dotted line) used to minimize model deviance and define the most parsimonious model. **C)** LASSO-derived probability scores stratify IpcPH and CpcPH patients, revealing significantly higher scores in CpcPH (****P < 0.0001). (**D–I**) transcriptomic expression of inflammatory markers in IpcPH and CpcPH animal lungs. **J)** Integrated Discrimination Improvement (IDI) analysis comparing NT-proBNP alone vs. NT-proBNP + protein biomarkers or LASSO score. Bars represent the improvement in predicted risk; asterisks denote significant enhancement ($p < 0.05$). **K)** Net Reclassification Index (NRI) quantifying correct risk reclassification when each marker or score is added to NT-proBNP. **L)** C-statistic comparison demonstrating prognostic discrimination of models combining NT-proBNP with individual biomarkers and LASSO score. Error bars represent 95% confidence intervals; asterisks indicate significant improvement. All survival analyses were assessed using log-rank tests. Data are expressed as mean ± SEM. Statistical comparisons were performed using unpaired t-test.

**Figure S6. Longitudinal echocardiographic evaluation of cardiac function in preclinical IpcPH and CpcPH models with or without levosimendan treatment.** Panels **A–G** show right and left ventricular parameters in IpcPH animals compared to Sham and IpcPH + Levosimendan groups: **A)** pulmonary artery acceleration time (PAAT), **B)** tricuspid annular plane systolic excursion (TAPSE), **C)** RV S′ wave, **D)** mitral annular plane systolic excursion (MAPSE), **E)** LV systolic velocity (LV-S′, E), **F)** E/E′ ratio, and **G)** body weight. Panels **H–N** show corresponding parameters in CpcPH animals versus Sham-Mets and CpcPH + Levosimendan groups. Data are expressed as mean ± SEM. Statistical comparisons were performed using repeated measures ANOVA followed by post hoc testing. *P < 0.05, **P < 0.01, ***P < 0.001, ****P < 0.0001 vs. respective controls.

**Figure S7. Histological and immunofluorescence assessment of left and right ventricular remodeling in IpcPH and CpcPH animals with or without levosimendan treatment. Panels A–D**

show left ventricular (LV) remodeling parameters including **A)** cardiomyocyte cross-sectional area (H&E), **B)** fibrosis (Masson's trichrome), **C)** $CD68^+$ macrophage infiltration, and **D)** $CD45^+$ leukocyte infiltration. **Panels E–H** present corresponding analyses in the right ventricle (RV): **E)** cardiomyocyte area (H&E), **F)** fibrosis (F, Masson's trichrome), **G)** $CD68^+$, and **H)** $CD45^+$ cell infiltration. Quantifications are shown as mean ± SEM and statistical comparisons were performed using one-way ANOVA followed by Tukey's post hoc test. ns = not significant, ** $P < 0.01$, **** $P < 0.0001$.

**Figure S8. Principal component analysis (PCA) of lung transcriptomic profiles in IpcPH animals with or without levosimendan treatment.** PCA plot shows that IpcPH and IpcPH + Levosimendan samples cluster closely with Sham animals, indicating minimal transcriptomic differences across these groups. This supports the lack of significant differential expression observed in IpcPH lungs with or without treatment.

**Figure S9. Transcriptomic alterations in the lungs of CpcPH animals relative to Sham-Mets controls**. **A)** Volcano plot showing differentially expressed genes (DEGs) in CpcPH versus Sham-Mets, with upregulated genes (red) and downregulated genes (green). (B–C) Functional enrichment analysis of **B)** downregulated and **C)** upregulated DEGs in CpcPH lungs, categorized by biological process (blue), cellular component (green), molecular function (purple), and KEGG pathways (orange). Enrichment score reflects the strength of association between gene sets and the corresponding functional category.

**Figure S10. Transcriptomic changes in the lungs of CpcPH animals following Levosimendan treatment. A)** Volcano plot comparing gene expression profiles between CpcPH and CpcPH + Levosimendan lungs. Differentially expressed genes (DEGs) upregulated in CpcPH + Levo are shown in red; downregulated DEGs are shown in green. Functional enrichment analysis of **B)** downregulated and **C)** upregulated DEGs following Levosimendan treatment, categorized by biological process (blue), cellular component (green), molecular function (purple), and KEGG pathways (orange). Results demonstrate a shift from inflammatory and immune-related gene signatures to metabolic pathways and fatty acid oxidation following treatment.

**Figure S11. Transcriptomic comparison between CpcPH animals treated with Levosimendan and Sham-Mets controls. A)** Volcano plot displaying differentially expressed genes (DEGs) between CpcPH + Levosimendan and Sham-Mets lungs. A small number of DEGs were identified, suggesting that Levosimendan treatment brings the lung transcriptomic profile of CpcPH animals closer to baseline. **B)** Heatmap illustrating expression levels of selected DEGs distinguishing CpcPH + Levo from Sham-Mets, including genes involved in circadian rhythm, inflammation, and extracellular matrix regulation. Color scale indicates relative gene expression (Z-score).

**Figure S12. Gene ontology analysis of differentially expressed genes (DEGs) reversed by Levosimendan in CpcPH lungs. A)** Quadrant plot comparing Log2 fold changes between CpcPH vs. Sham-Mets (x-axis) and CpcPH + Levo vs. CpcPH (y-axis) lungs. Genes upregulated in CpcPH and downregulated by Levosimendan (n=137, blue quadrant), and genes downregulated in CpcPH and upregulated by Levosimendan (n=63, red quadrant), were subjected to pathway enrichment. **B)** Enriched pathways for the 63 downregulated genes reversed by Levosimendan, revealing alterations in NADPH oxidase activity, fatty acid metabolism, and oxidative stress pathways. **C)** Enriched pathways for the 137 upregulated genes reversed by Levosimendan, highlighting attenuation of inflammatory responses, cytokine signaling, and extracellular matrix remodeling.

**Figure S13. Levosimendan treatment attenuates NF-κB signaling in the lungs of CpcPH rats.**
**A)** Transcription factor enrichment analysis of genes upregulated in CpcPH and downregulated following levosimendan treatment (CpcPH↑/Levo↓ DEGs), and of all Levo↓ DEGs. NF-κB emerged as the top enriched transcription factor. **B)** Western blot analysis of phosphorylated NF-κB (p-NF-κB) in lung tissue from CpcPH animals treated with levosimendan. Levosimendan significantly reduced NF-κB activation. Data are presented as mean ± SEM. Statistical comparisons were performed using one-way repeated measures ANOVA followed by post hoc testing. *$P < 0.05$ vs. respective controls.

**Figure S14. Levosimendan treatment reduces the expression of pro-inflammatory markers in the lungs of CpcPH animals. A)** Representative immunofluorescence images illustrate diminished staining intensity in distal pulmonary arteries of levosimendan-treated animals compared to controls. (**B–G**) Proteomic quantification of inflammatory markers in lung tissue of levosimendan treated animals. Data are presented as mean ± SEM. Statistical comparisons were performed using one-way repeated measures ANOVA followed by post hoc testing. *P < 0.05, **P < 0.01 , ***P < 0.005, ****P < 0.0005 vs. respective controls.

A)

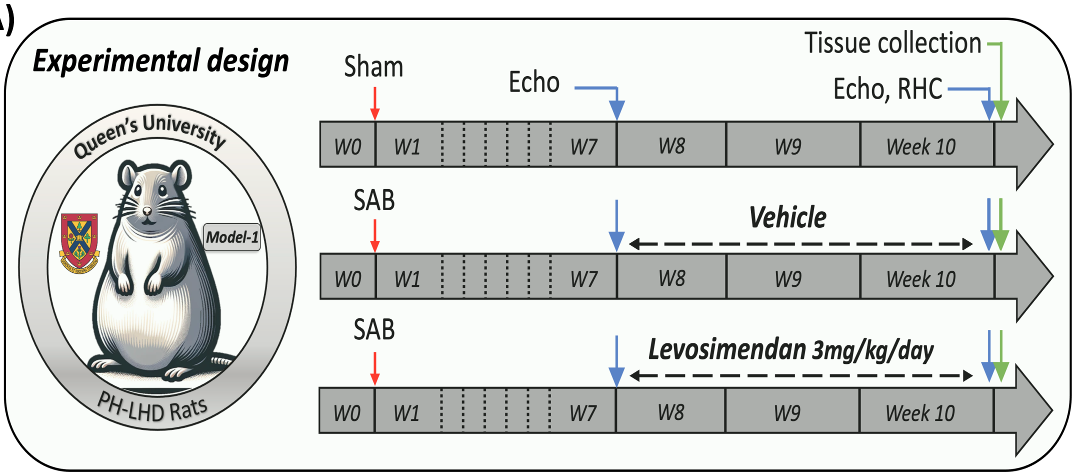
Experimental design
Queen's University
Model-1
PH-LHD Rats
Sham
Echo
Tissue collection
Echo, RHC
W0
W1
W7
W8
W9
Week 10
SAB
Vehicle
SAB
Levosimendan 3mg/kg/day

B)

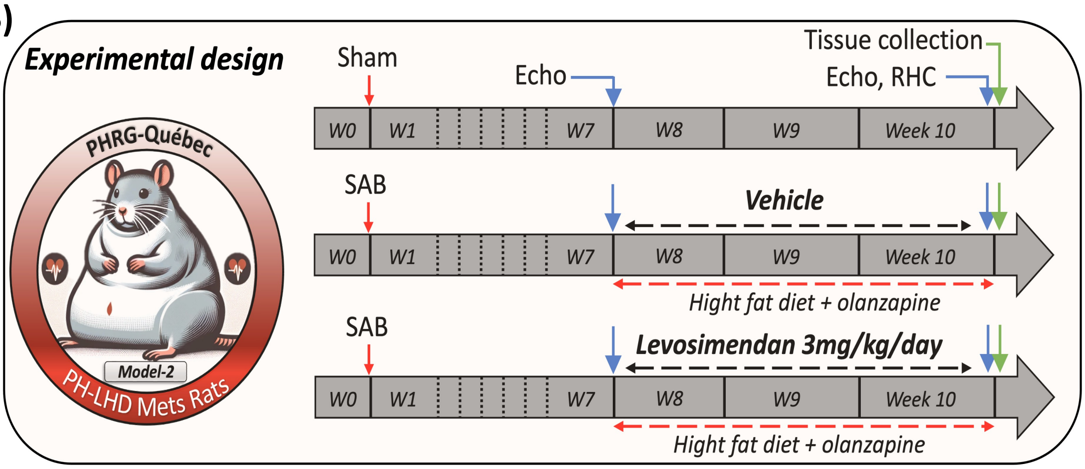
Experimental design
PHRG-Québec
Model-2
PH-LHD Mets Rats
Sham
Echo
Tissue collection
Echo, RHC
W0
W1
W7
W8
W9
Week 10
SAB
Vehicle
Hight fat diet + olanzapine
SAB
Levosimendan 3mg/kg/day
Hight fat diet + olanzapine

**A)**

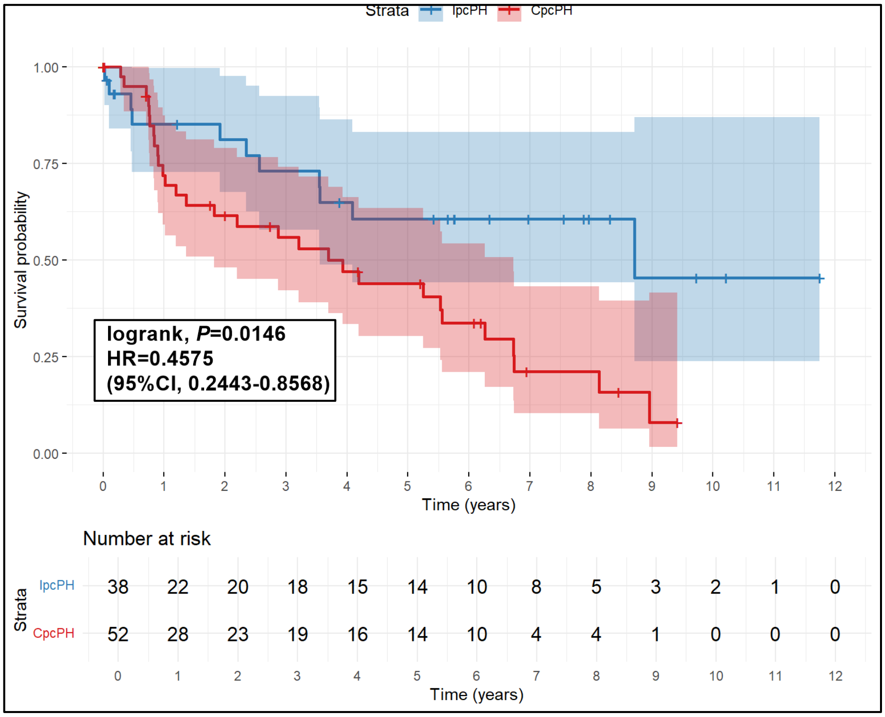

Strata
IpcPH
CpcPH
Survival probability
1.00
0.75
0.50
0.25
0.00
logrank, *P*=0.0146
HR=0.4575
(95%CI, 0.2443-0.8568)
0 1 2 3 4 5 6 7 8 9 10 11 12
Time (years)
Number at risk
Strata
IpcPH 38 22 20 18 15 14 10 8 5 3 2 1 0
CpcPH 52 28 23 19 16 14 10 4 4 1 0 0 0
0 1 2 3 4 5 6 7 8 9 10 11 12
Time (years)

A)

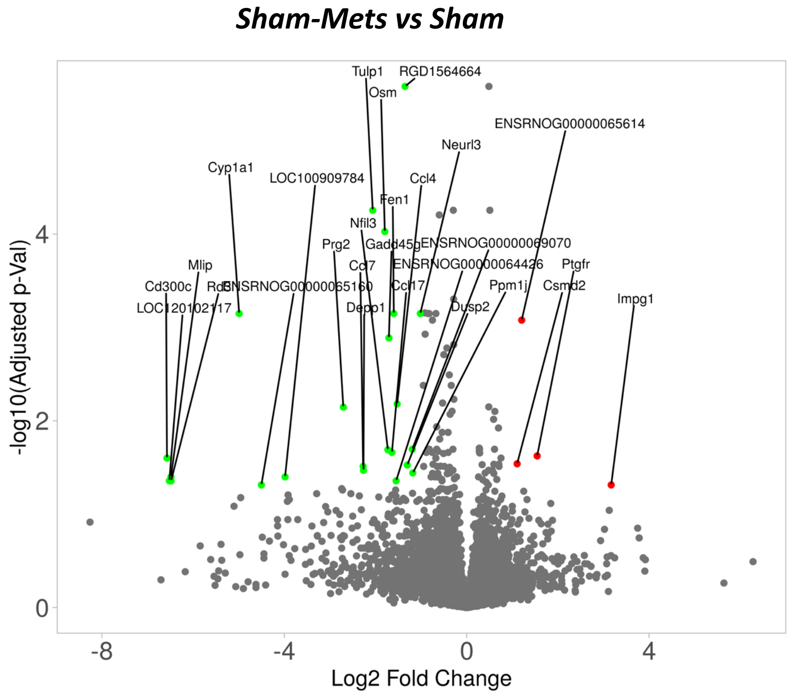

Sham-Mets vs Sham
Tulp1
RGD1564664
Osm
ENSRNOG00000065614
Neurl3
Cyp1a1
LOC100909784
Ccl4
Fen1
Nfil3
Prg2
Gadd45g
ENSRNOG00000069070
Mlip
Ccl7
ENSRNOG00000064426
Ptgfr
Cd300c
Rd3l
ENSRNOG00000065160
Ccl17
Ppm1j
Csmd2
Impg1
LOC120102117
Depp1
Dusp2
-log10(Adjusted p-Val)
Log2 Fold Change
-8
-4
0
4
2


B)

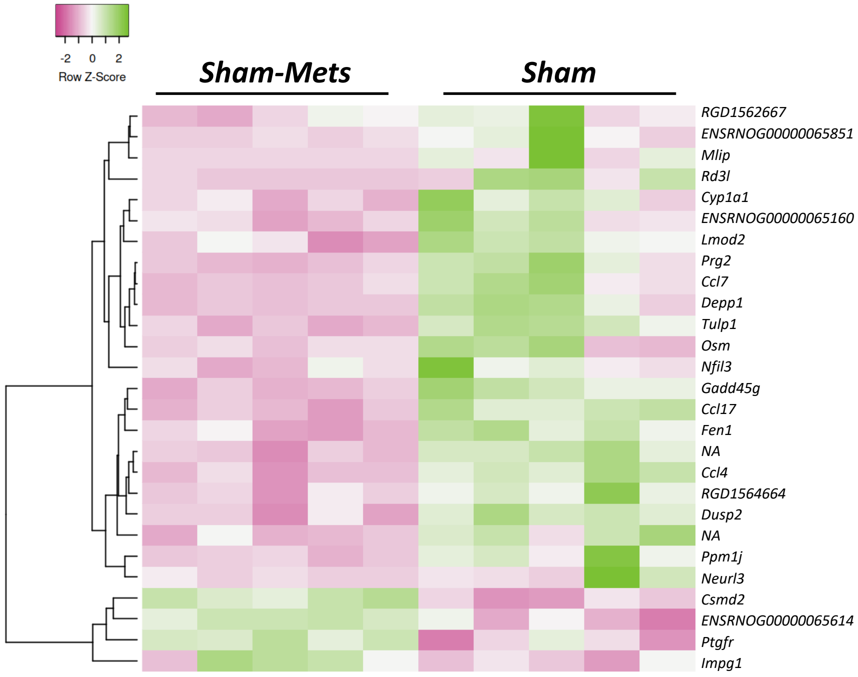

-2 0 2
Row Z-Score
Sham-Mets
Sham
RGD1562667
ENSRNOG00000065851
Mlip
Rd3l
Cyp1a1
ENSRNOG00000065160
Lmod2
Prg2
Ccl7
Depp1
Tulp1
Osm
Nfil3
Gadd45g
Ccl17
Fen1
NA
Ccl4
RGD1564664
Dusp2
NA
Ppm1j
Neurl3
Csmd2
ENSRNOG00000065614
Ptgfr
Impg1

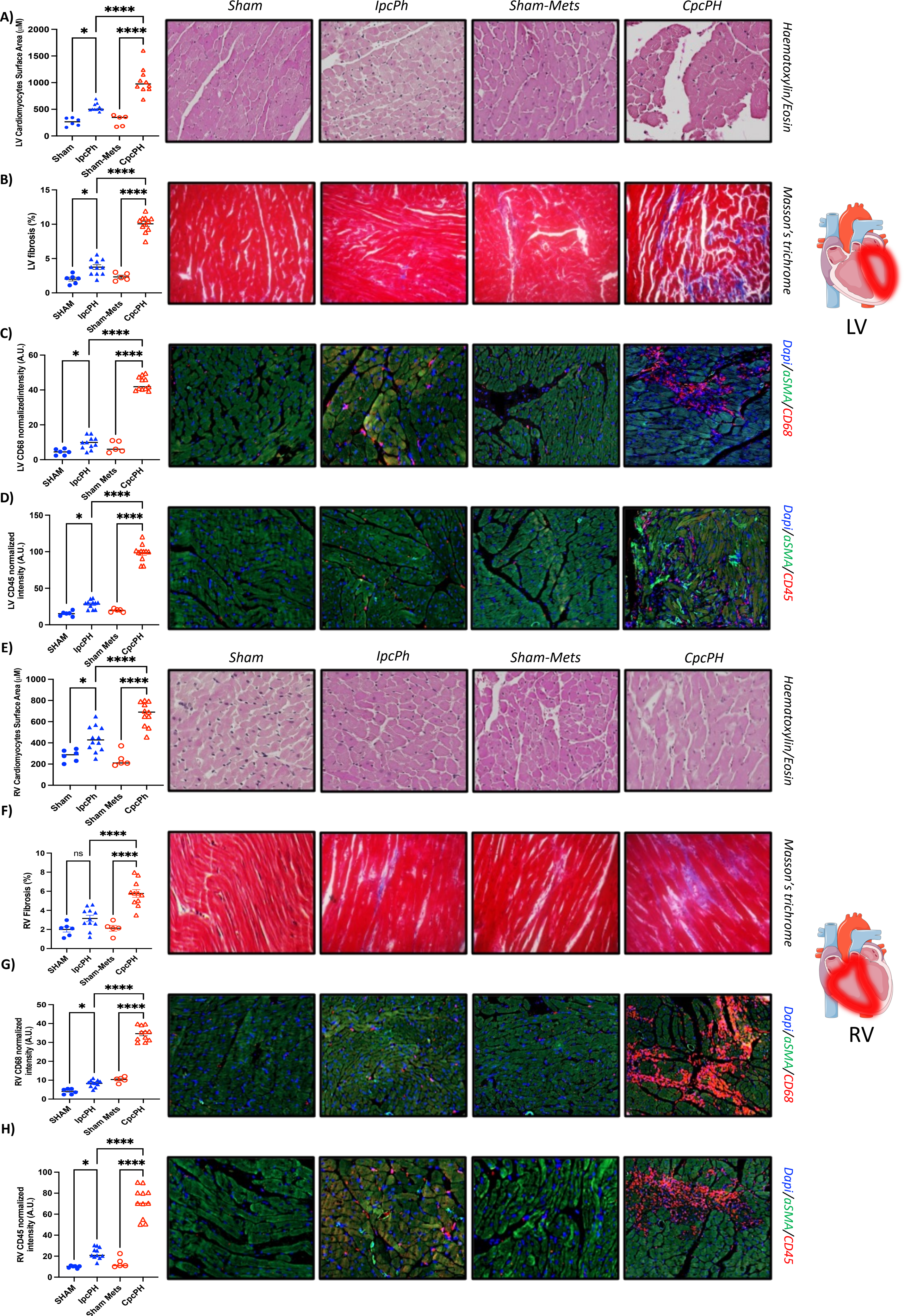

A)
B)
C)
D)
E)
F)
G)
H)
Sham
IpcPh
Sham-Mets
CpcPH
Haematoxylin/Eosin
Masson's trichrome
Dapi/aSMA/CD68
Dapi/aSMA/CD45
LV Cardiomyocytes Surface Area (μM)
LV fibrosis (%)
LV CD68 normalizedintensity (A.U.)
LV CD45 normalized intensity (A.U.)
RV Cardiomyocytes Surface Area (μM)
RV Fibrosis (%)
RV CD68 normalized intensity (A.U.)
RV CD45 normalized intensity (A.U.)
SHAM
IpcPH
Sham-Mets
Sham Mets
CpcPH
ns
LV
RV

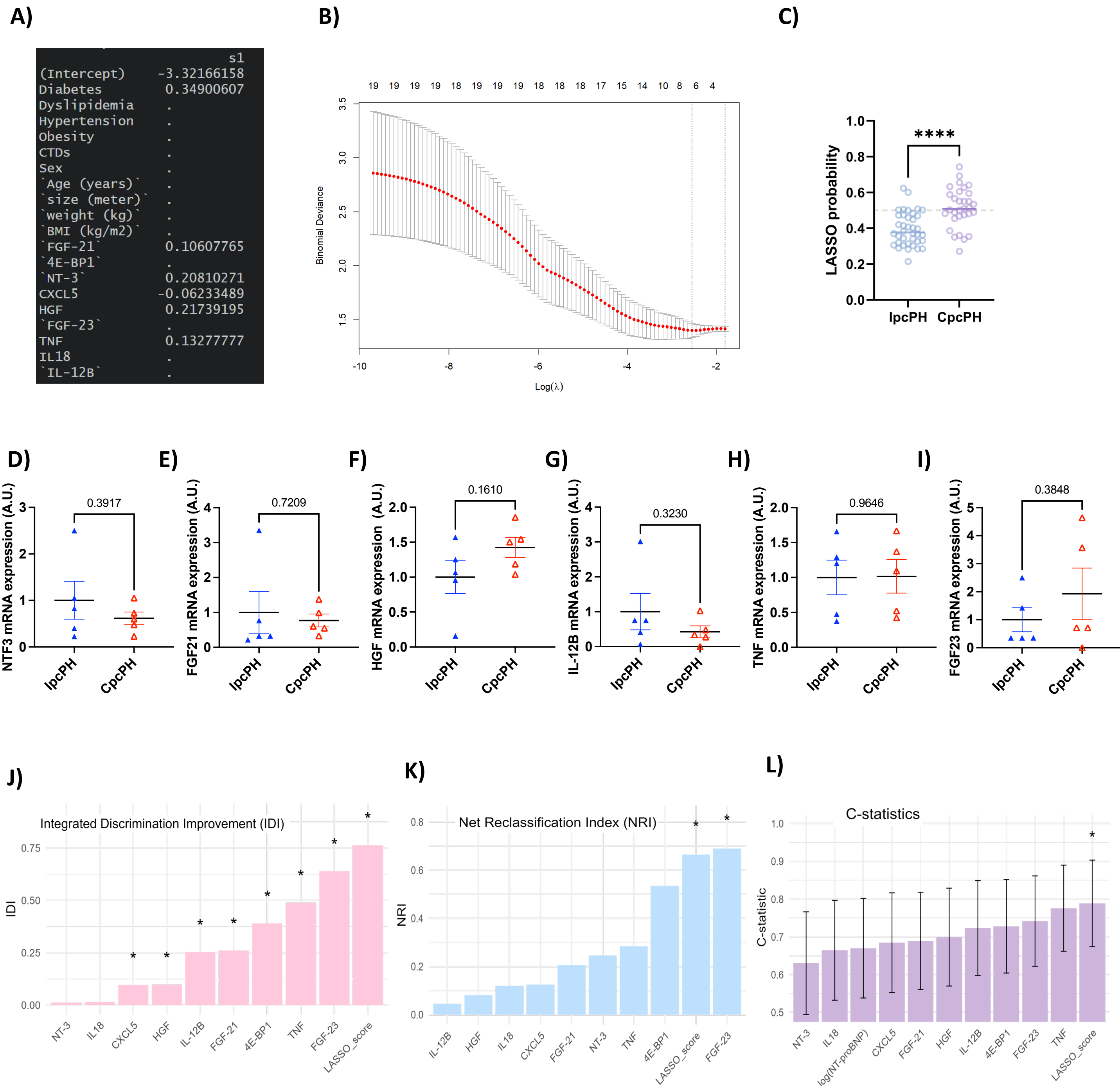
A)
s1
(Intercept) -3.32166158
Diabetes 0.34900607
Dyslipidemia .
Hypertension .
Obesity .
CTDs .
Sex .
`Age (years)` .
`size (meter)` .
`weight (kg)` .
`BMI (kg/m2)` .
`FGF-21` 0.10607765
`4E-BP1` .
`NT-3` 0.20810271
CXCL5 -0.06233489
HGF 0.21739195
`FGF-23` .
TNF 0.13277777
IL18 .
`IL-12B` .
B)
Binomial Deviance
Log(λ)
C)
LASSO probability
****
IpcPH
CpcPH
D)
NTF3 mRNA expression (A.U.)
0.3917
E)
FGF21 mRNA expression (A.U.)
0.7209
F)
HGF mRNA expression (A.U.)
0.1610
G)
IL-12B mRNA expression (A.U.)
0.3230
H)
TNF mRNA expression (A.U.)
0.9646
I)
FGF23 mRNA expression (A.U.)
0.3848
J)
Integrated Discrimination Improvement (IDI)
IDI
NT-3
IL18
CXCL5
HGF
IL-12B
FGF-21
4E-BP1
TNF
FGF-23
LASSO_score
K)
Net Reclassification Index (NRI)
NRI
IL-12B
HGF
IL18
CXCL5
FGF-21
NT-3
TNF
4E-BP1
LASSO_score
FGF-23
L)
C-statistics
C-statistic
NT-3
IL18
log(NT-proBNP)
CXCL5
FGF-21
HGF
IL-12B
4E-BP1
FGF-23
TNF
LASSO_score

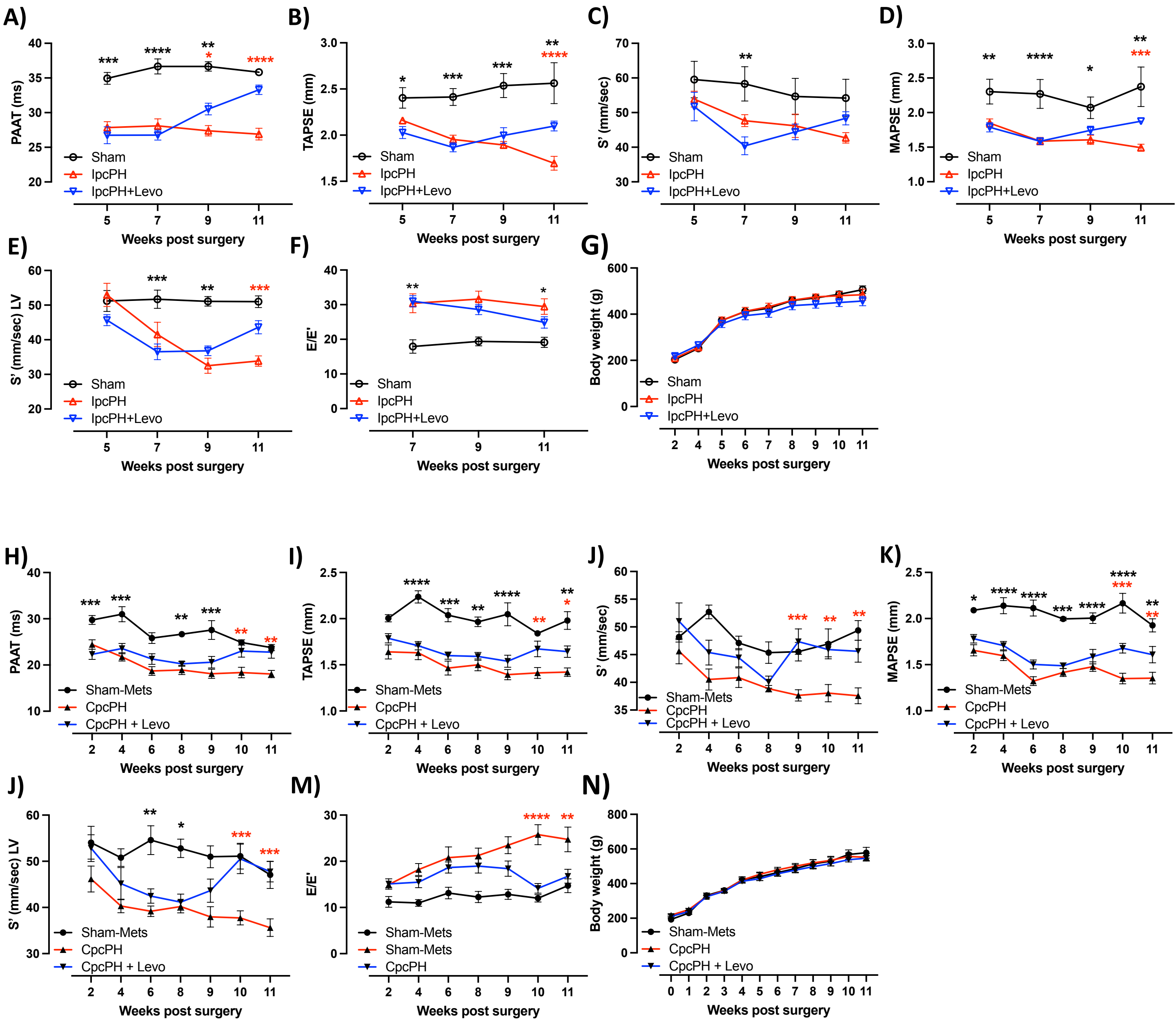

A)
PAAT (ms)
Sham
IpcPH
IpcPH+Levo
Weeks post surgery
B)
TAPSE (mm)
C)
S' (mm/sec)
D)
MAPSE (mm)
E)
S' (mm/sec) LV
F)
E/E'
G)
Body weight (g)
H)
Sham-Mets
CpcPH
CpcPH + Levo
I)
J)
K)
J)
M)
Sham-Mets
Sham-Mets
CpcPH
N)

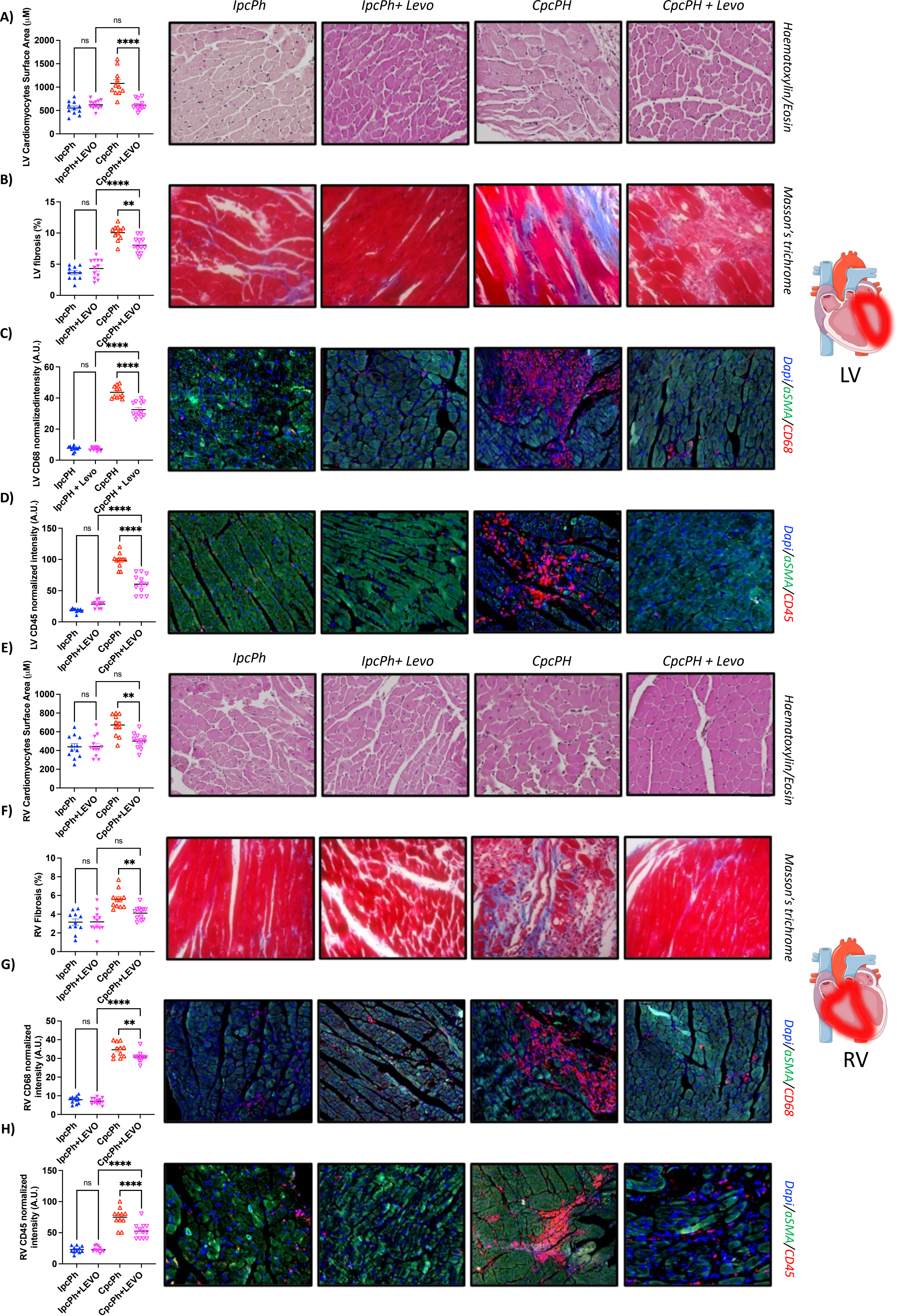
A)
B)
C)
D)
E)
F)
G)
H)
IpcPh
IpcPh+ Levo
CpcPH
CpcPH + Levo
Haematoxylin/Eosin
Masson's trichrome
Dapi/αSMA/CD68
Dapi/αSMA/CD45
LV
RV
LV Cardiomyocytes Surface Area (μM)
LV fibrosis (%)
LV CD68 normalizedintensity (A.U.)
LV CD45 normalized intensity (A.U.)
RV Cardiomyocytes Surface Area (μM)
RV Fibrosis (%)
RV CD68 normalized intensity (A.U.)
RV CD45 normalized intensity (A.U.)
IpcPh
IpcPh+LEVO
CpcPh
CpcPh+LEVO
ns
****
**

**A)**

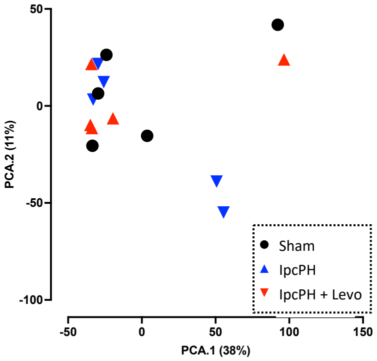

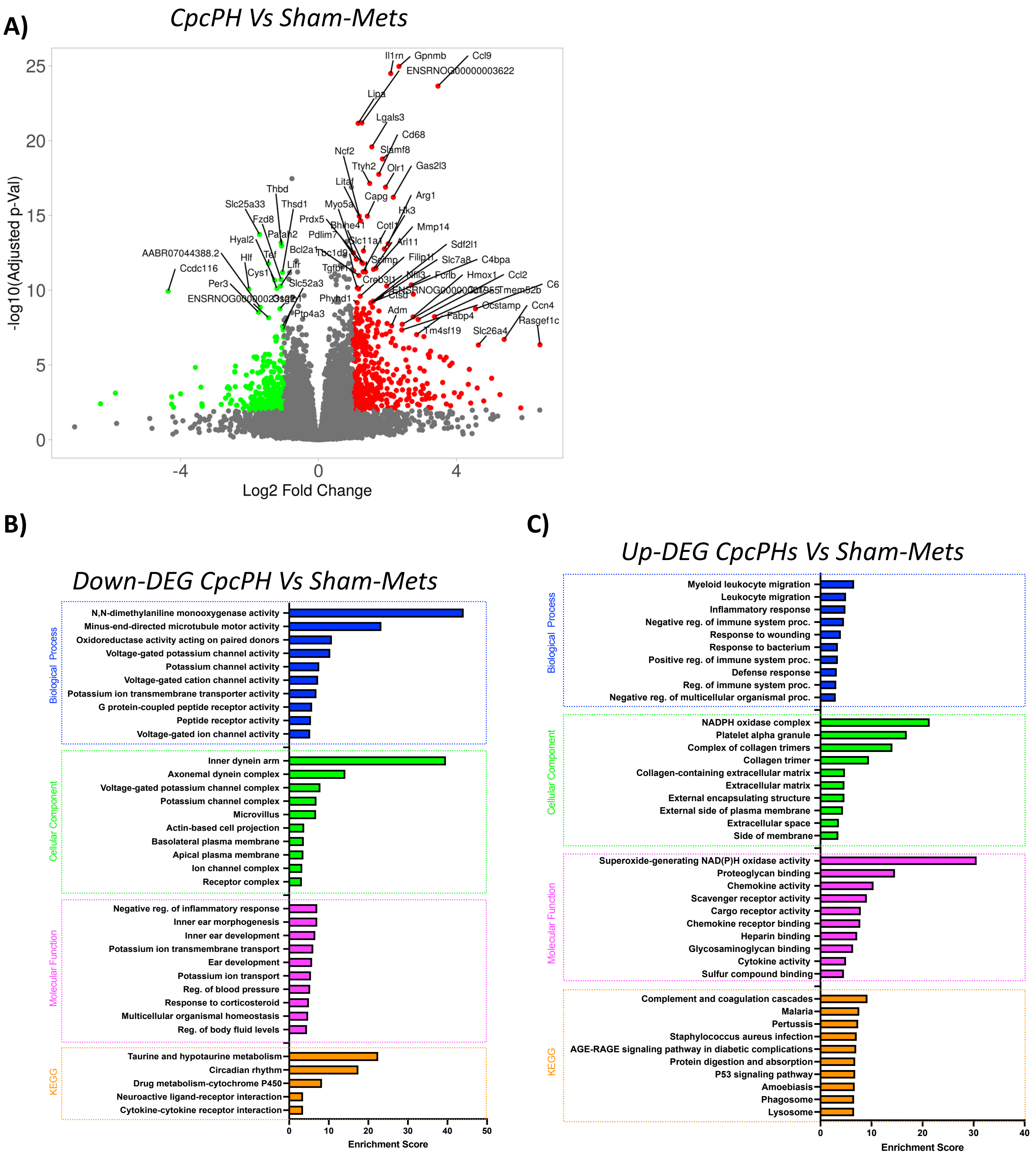

A)
CpcPH Vs Sham-Mets
-log10(Adjusted p-Val)
Log2 Fold Change
Il1rn
Gpnmb
Ccl9
ENSRNOG00000003622
Lipa
Lgals3
Cd68
Ncf2
Slamf8
Ttyh2
Olr1
Gas2l3
Litaf
Capg
Arg1
Thbd
Slc25a33
Thsd1
Myo5a
Hk3
Fzd8
Prdx5
Bhlhe41
Cotl1
Mmp14
Hyal2
Pafah2
Pdlim7
Slc11a1
Arl11
Sdf2l1
AABR07044388.2
Hlf
Tef
Bcl2a1
Tbc1d9
Scimp
Filip1l
Slc7a8
C4bpa
Ccdc116
Cys1
Lifr
Tgfbr1
Creb3l1
Nfil3
Fcrlb
Hmox1
Ccl2
Per3
Slc52a3
C6
ENSRNOG00000023921
Ctsd
Phyhd1
Tmem52b
Ptp4a3
Adm
Ocstamp
Ccn4
Fabp4
Rasgef1c
Tm4sf19
Slc26a4
B)
Down-DEG CpcPH Vs Sham-Mets
Biological Process
N,N-dimethylaniline monooxygenase activity
Minus-end-directed microtubule motor activity
Oxidoreductase activity acting on paired donors
Voltage-gated potassium channel activity
Potassium channel activity
Voltage-gated cation channel activity
Potassium ion transmembrane transporter activity
G protein-coupled peptide receptor activity
Peptide receptor activity
Voltage-gated ion channel activity
Cellular Component
Inner dynein arm
Axonemal dynein complex
Voltage-gated potassium channel complex
Potassium channel complex
Microvillus
Actin-based cell projection
Basolateral plasma membrane
Apical plasma membrane
Ion channel complex
Receptor complex
Molecular Function
Negative reg. of inflammatory response
Inner ear morphogenesis
Inner ear development
Potassium ion transmembrane transport
Ear development
Potassium ion transport
Reg. of blood pressure
Response to corticosteroid
Multicellular organismal homeostasis
Reg. of body fluid levels
KEGG
Taurine and hypotaurine metabolism
Circadian rhythm
Drug metabolism-cytochrome P450
Neuroactive ligand-receptor interaction
Cytokine-cytokine receptor interaction
Enrichment Score
C)
Up-DEG CpcPHs Vs Sham-Mets
Biological Process
Myeloid leukocyte migration
Leukocyte migration
Inflammatory response
Negative reg. of immune system proc.
Response to wounding
Response to bacterium
Positive reg. of immune system proc.
Defense response
Reg. of immune system proc.
Negative reg. of multicellular organismal proc.
Cellular Component
NADPH oxidase complex
Platelet alpha granule
Complex of collagen trimers
Collagen trimer
Collagen-containing extracellular matrix
Extracellular matrix
External encapsulating structure
External side of plasma membrane
Extracellular space
Side of membrane
Molecular Function
Superoxide-generating NAD(P)H oxidase activity
Proteoglycan binding
Chemokine activity
Scavenger receptor activity
Cargo receptor activity
Chemokine receptor binding
Heparin binding
Glycosaminoglycan binding
Cytokine activity
Sulfur compound binding
KEGG
Complement and coagulation cascades
Malaria
Pertussis
Staphylococcus aureus infection
AGE-RAGE signaling pathway in diabetic complications
Protein digestion and absorption
P53 signaling pathway
Amoebiasis
Phagosome
Lysosome
Enrichment Score

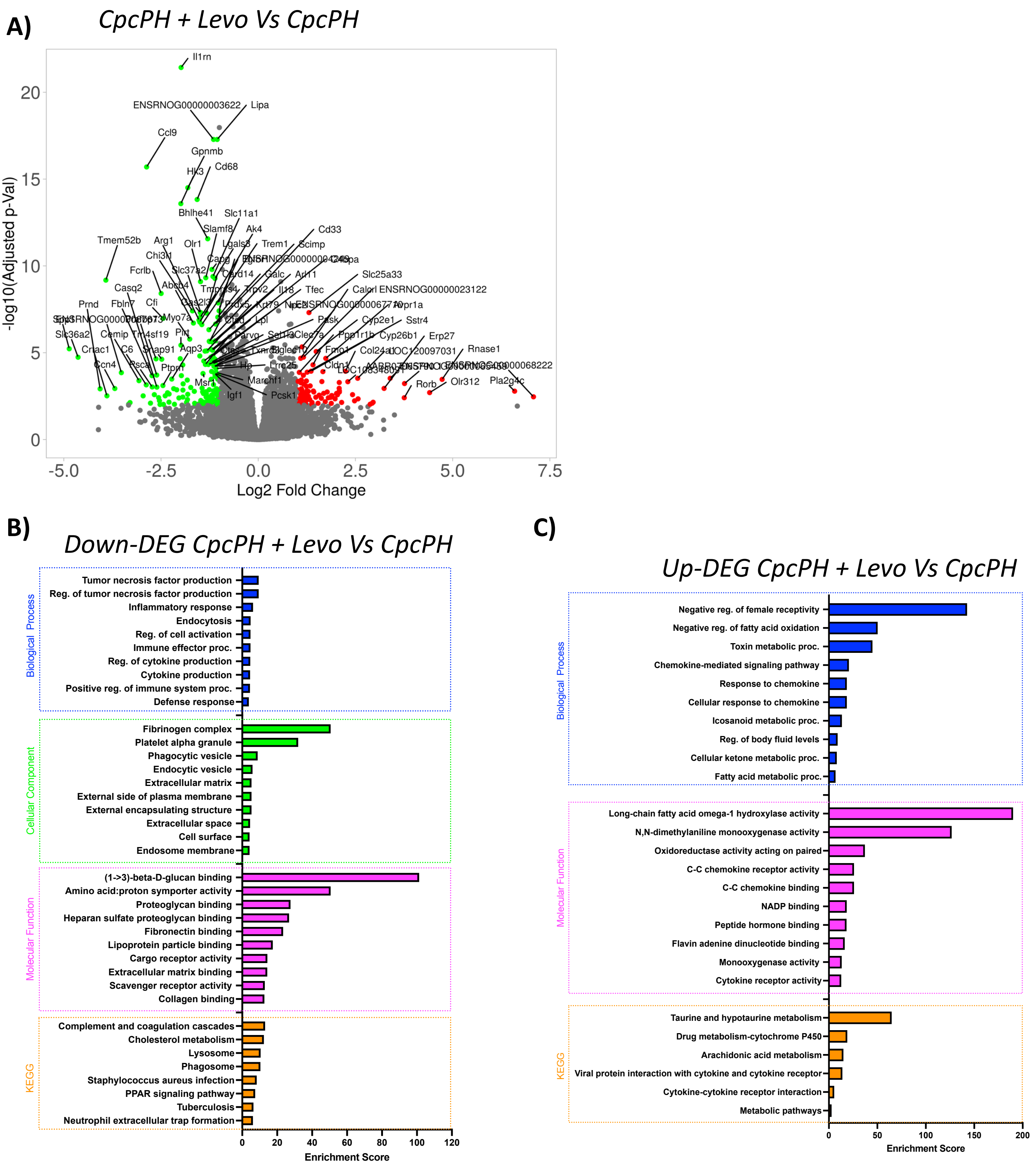
A)
CpcPH + Levo Vs CpcPH
-log10(Adjusted p-Val)
Log2 Fold Change
-5.0
-2.5
0.0
2.5
5.0
7.5
0
5
10
15
20
Il1rn
ENSRNOG00000003622
Lipa
Ccl9
Gpnmb
Hk3
Cd68
Bhlhe41
Slc11a1
Slamf8
Ak4
Tmem52b
Arg1
Olr1
Lgals3
Trem1
Scimp
Cd33
Chi3l1
Fcrlb
Slc37a2
Card14
Galc
Arl11
Slc25a33
Casq2
Abcb4
Calcrl
ENSRNOG00000023122
Prnd
Fbln7
Cfi
Myo7a
Lpl
Pask
Cyp2e1
Sstr4
Slc36a2
Cemip
Tm4sf19
Plxt
Ppp1r1b
Cyp26b1
Erp27
Crlac1
C6
Snap91
Aqp3
Fmo1
Col24a1
LOC120097031
Rnase1
Ccn4
Psca
Ptprn
Hp
Lrrc25
Cldn1
ENSRNOG00000068222
Msr1
March1
Rorb
Olr312
Pla2g4c
Igf1
Pcsk1
B)
Down-DEG CpcPH + Levo Vs CpcPH
Biological Process
Tumor necrosis factor production
Reg. of tumor necrosis factor production
Inflammatory response
Endocytosis
Reg. of cell activation
Immune effector proc.
Reg. of cytokine production
Cytokine production
Positive reg. of immune system proc.
Defense response
Cellular Component
Fibrinogen complex
Platelet alpha granule
Phagocytic vesicle
Endocytic vesicle
Extracellular matrix
External side of plasma membrane
External encapsulating structure
Extracellular space
Cell surface
Endosome membrane
Molecular Function
(1->3)-beta-D-glucan binding
Amino acid:proton symporter activity
Proteoglycan binding
Heparan sulfate proteoglycan binding
Fibronectin binding
Lipoprotein particle binding
Cargo receptor activity
Extracellular matrix binding
Scavenger receptor activity
Collagen binding
KEGG
Complement and coagulation cascades
Cholesterol metabolism
Lysosome
Phagosome
Staphylococcus aureus infection
PPAR signaling pathway
Tuberculosis
Neutrophil extracellular trap formation
0
20
40
60
80
100
120
Enrichment Score
C)
Up-DEG CpcPH + Levo Vs CpcPH
Biological Process
Negative reg. of female receptivity
Negative reg. of fatty acid oxidation
Toxin metabolic proc.
Chemokine-mediated signaling pathway
Response to chemokine
Cellular response to chemokine
Icosanoid metabolic proc.
Reg. of body fluid levels
Cellular ketone metabolic proc.
Fatty acid metabolic proc.
Molecular Function
Long-chain fatty acid omega-1 hydroxylase activity
N,N-dimethylaniline monooxygenase activity
Oxidoreductase activity acting on paired
C-C chemokine receptor activity
C-C chemokine binding
NADP binding
Peptide hormone binding
Flavin adenine dinucleotide binding
Monooxygenase activity
Cytokine receptor activity
KEGG
Taurine and hypotaurine metabolism
Drug metabolism-cytochrome P450
Arachidonic acid metabolism
Viral protein interaction with cytokine and cytokine receptor
Cytokine-cytokine receptor interaction
Metabolic pathways
0
50
100
150
200
Enrichment Score

A)

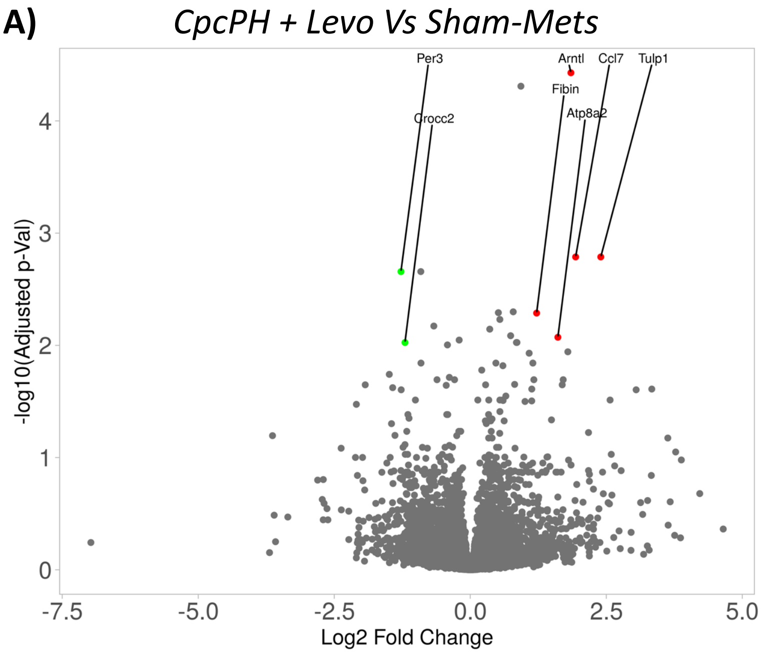

CpcPH + Levo Vs Sham-Mets
Per3
Crocc2
Arntl
Ccl7
Tulp1
Fibin
Atp8a2
-log10(Adjusted p-Val)
Log2 Fold Change
-7.5
-5.0
-2.5
0.0
2.5
5.0
0
1
2
3
4


B)

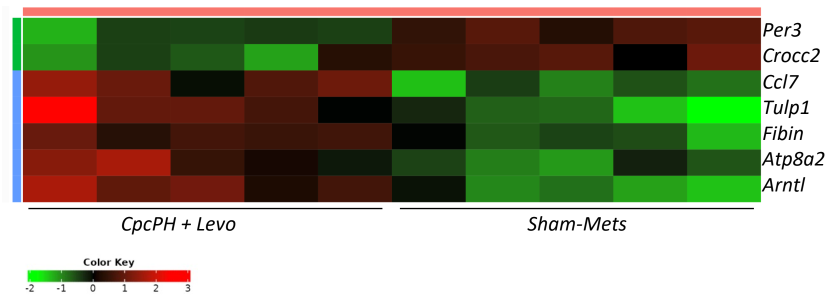

Per3
Crocc2
Ccl7
Tulp1
Fibin
Atp8a2
Arntl
CpcPH + Levo
Sham-Mets
Color Key
-2
-1
0
1
2
3

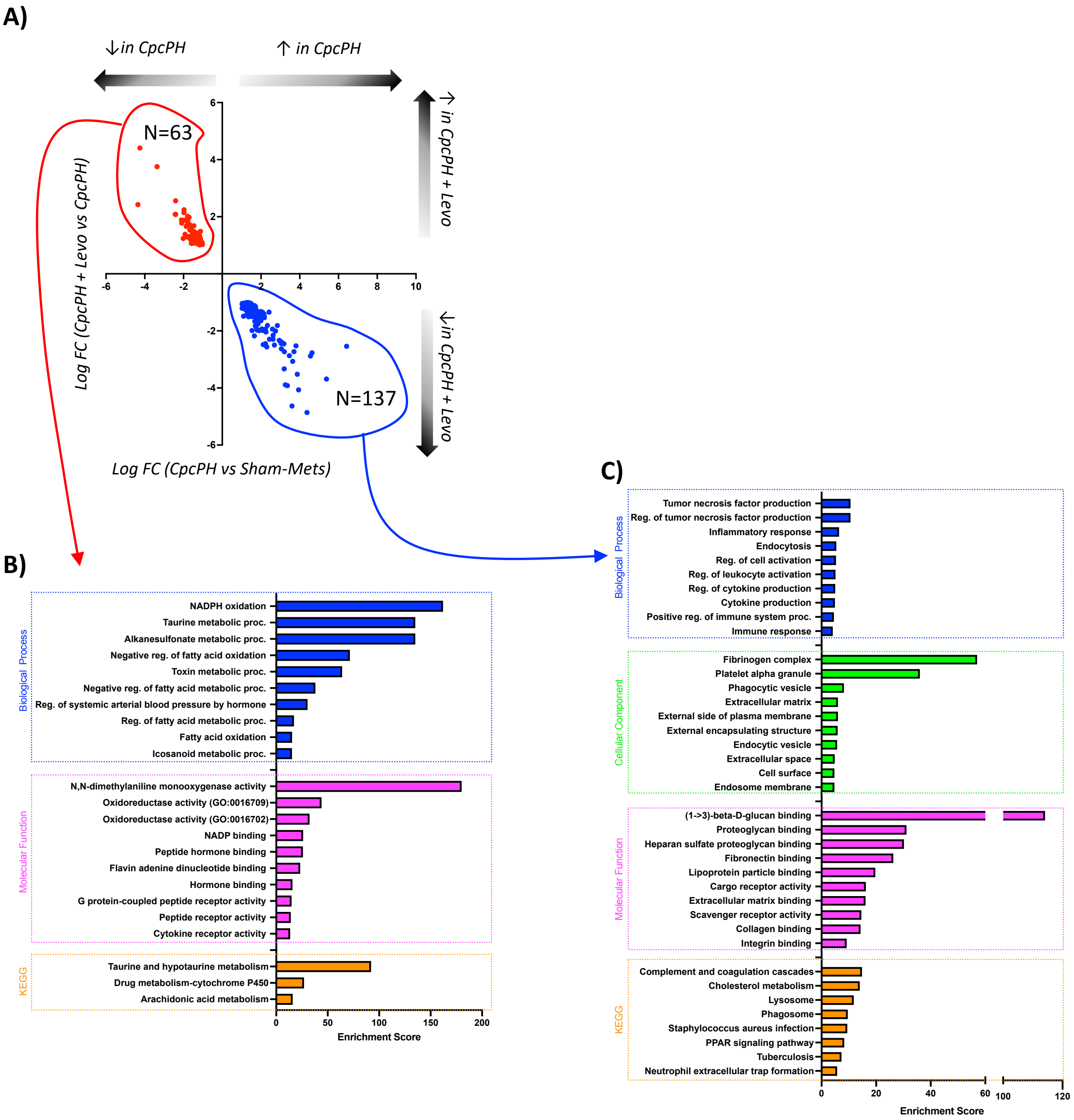
A)
↓in CpcPH
↑ in CpcPH
↑ in CpcPH + Levo
↓in CpcPH + Levo
N=63
N=137
Log FC (CpcPH + Levo vs CpcPH)
Log FC (CpcPH vs Sham-Mets)
B)
Biological Process
NADPH oxidation
Taurine metabolic proc.
Alkanesulfonate metabolic proc.
Negative reg. of fatty acid oxidation
Toxin metabolic proc.
Negative reg. of fatty acid metabolic proc.
Reg. of systemic arterial blood pressure by hormone
Reg. of fatty acid metabolic proc.
Fatty acid oxidation
Icosanoid metabolic proc.
Molecular Function
N,N-dimethylaniline monooxygenase activity
Oxidoreductase activity (GO:0016709)
Oxidoreductase activity (GO:0016702)
NADP binding
Peptide hormone binding
Flavin adenine dinucleotide binding
Hormone binding
G protein-coupled peptide receptor activity
Peptide receptor activity
Cytokine receptor activity
KEGG
Taurine and hypotaurine metabolism
Drug metabolism-cytochrome P450
Arachidonic acid metabolism
Enrichment Score
C)
Biological Process
Tumor necrosis factor production
Reg. of tumor necrosis factor production
Inflammatory response
Endocytosis
Reg. of cell activation
Reg. of leukocyte activation
Reg. of cytokine production
Cytokine production
Positive reg. of immune system proc.
Immune response
Cellular Component
Fibrinogen complex
Platelet alpha granule
Phagocytic vesicle
Extracellular matrix
External side of plasma membrane
External encapsulating structure
Endocytic vesicle
Extracellular space
Cell surface
Endosome membrane
Molecular Function
(1->3)-beta-D-glucan binding
Proteoglycan binding
Heparan sulfate proteoglycan binding
Fibronectin binding
Lipoprotein particle binding
Cargo receptor activity
Extracellular matrix binding
Scavenger receptor activity
Collagen binding
Integrin binding
KEGG
Complement and coagulation cascades
Cholesterol metabolism
Lysosome
Phagosome
Staphylococcus aureus infection
PPAR signaling pathway
Tuberculosis
Neutrophil extracellular trap formation
Enrichment Score

A)

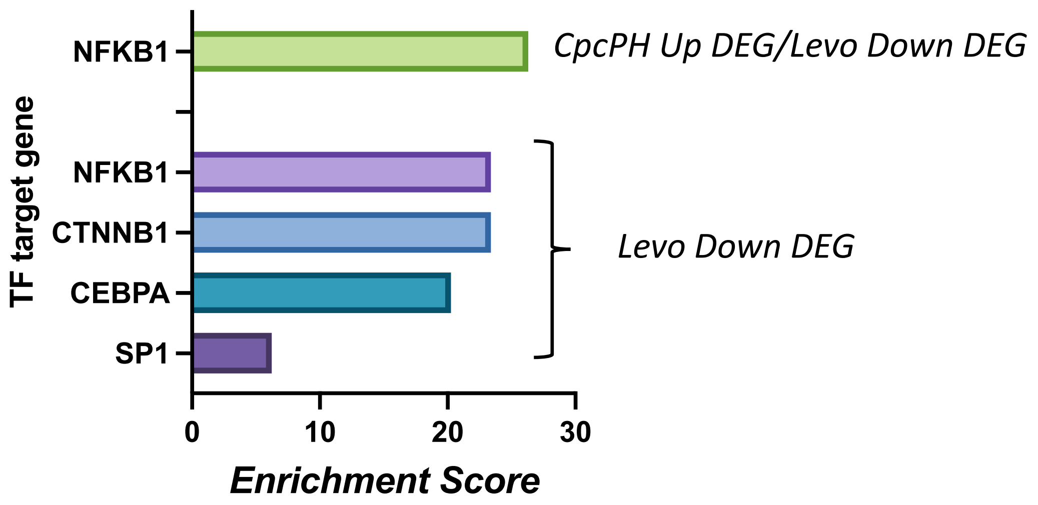
NFKB1
CpcPH Up DEG/Levo Down DEG
TF target gene
NFKB1
CTNNB1
CEBPA
SP1
Levo Down DEG
0
10
20
30
Enrichment Score

B)

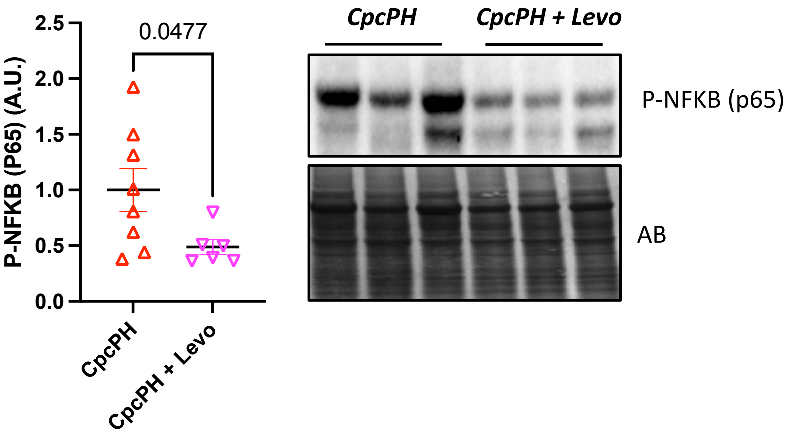
P-NFKB (P65) (A.U.)
2.5
2.0
1.5
1.0
0.5
0.0
0.0477
CpcPH
CpcPH + Levo
CpcPH
CpcPH + Levo
P-NFKB (p65)
AB

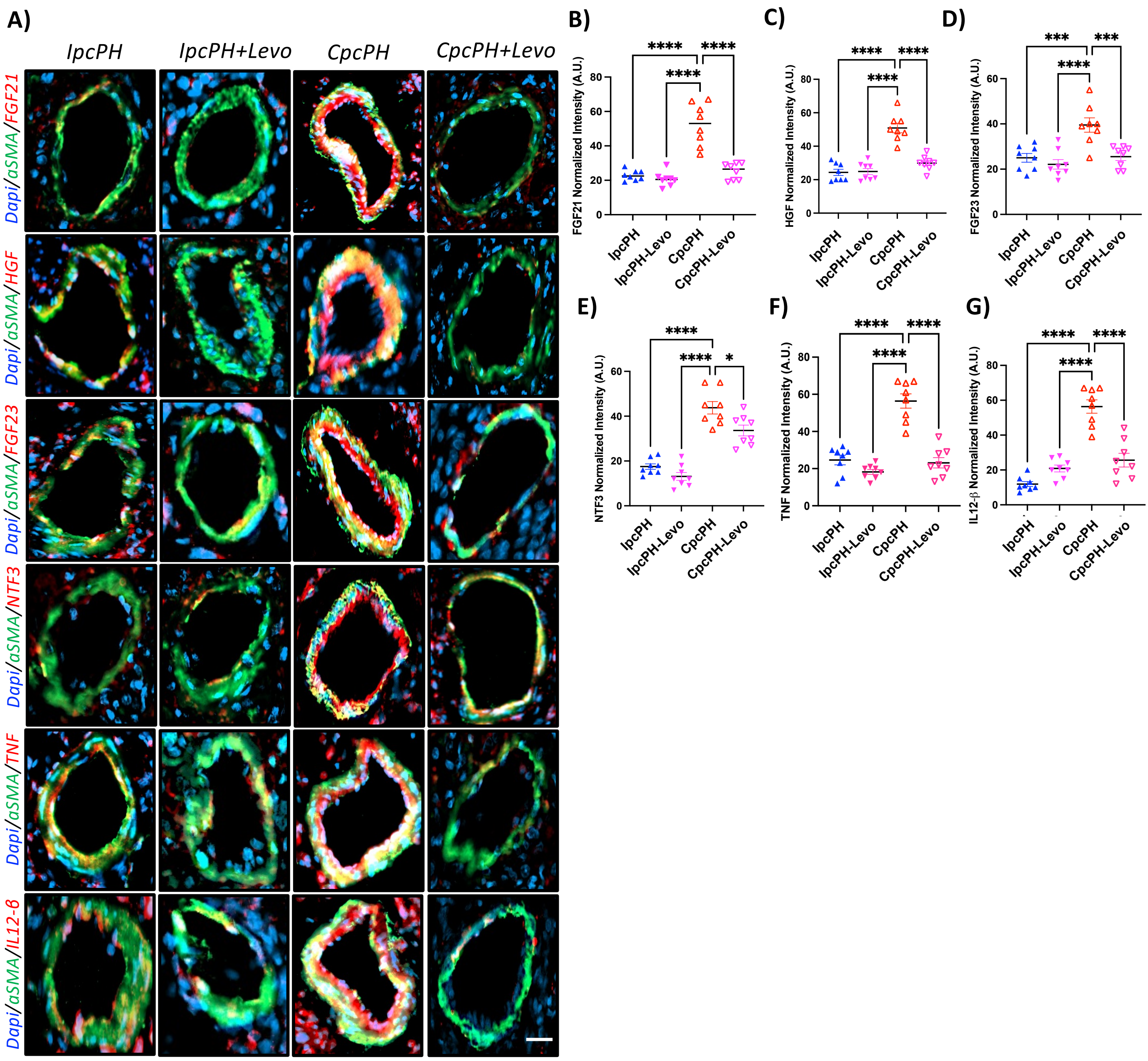

A)
IpcPH
IpcPH+Levo
CpcPH
CpcPH+Levo
Dapi/aSMA/FGF21
Dapi/aSMA/HGF
Dapi/aSMA/FGF23
Dapi/aSMA/NTF3
Dapi/aSMA/TNF
Dapi/aSMA/IL12-β
B)
FGF21 Normalized Intensity (A.U.)
C)
HGF Normalized Intensity (A.U.)
D)
FGF23 Normalized Intensity (A.U.)
E)
NTF3 Normalized Intensity (A.U.)
F)
TNF Normalized Intensity (A.U.)
G)
IL12-β Normalized Intensity (A.U.)
IpcPH
IpcPH-Levo
CpcPH
CpcPH-Levo
